# A MaBBX21-MaGCN5 Histone Acetyltransferase Module Regulates Flavonoid Biosynthesis to Improve Heat and UV-B Stress Tolerance in Banana (*Musa acuminata*)

**DOI:** 10.64898/2026.04.21.719834

**Authors:** Samar Singh, Shivi Tyagi, Ram Naresh, Sweta Bhambhani, Himani Chhatwal, Jogindra Naik, Boas Pucker, Ralf Stracke, Ashutosh Pandey

## Abstract

Flavonoids are important specialized metabolites that contribute to plant pigmentation, stress adaptation, and nutritional value. In banana, a major global staple crop, their accumulation is highly tissue-specific, with very low levels in the edible pulp, and the mechanisms underlying this spatial distribution remain unclear. Here, integrative transcriptomic and metabolomic analyses across vegetative and reproductive tissues reveal that light-responsive regulatory networks control tissue-specific flavonoid biosynthesis. We identified a B-box transcription factor MaBBX21 as a key positive regulator of flavonoid biosynthesis. Its overexpression enhances flavonoid accumulation, whereas knockdown leads to a reduction in flavonoid levels in banana. Mechanistically, MaBBX21 interacts with MaHY5 and directly activates anthocyanin biosynthesis genes (*MaDFR2* and *MaANS*). It also regulates metabolic flux by binding to the *MaWRKY23* promoter, promoting flavonol biosynthesis through activation of *MaFLS1* while partially repressing the anthocyanin branch. In addition, MaBBX21 introduces an epigenetic layer by activating the *histone acetyltransferase MaGCN5,* increasing H3K9 acetylation at target promoters, including *MaDFR2, MaANS,* and MaBBX21 itself, thereby forming a positive feedback loop. Functionally, *MaBBX21* overexpression enhances flavonoid accumulation, ROS scavenging, and tolerance to heat and UV-B stress, whereas knockdown lines show reduced metabolite levels and increased stress sensitivity. Collectively, these results define a MaBBX21–MaWRKY23/MaGCN5 regulatory axis that integrates transcriptional regulation, and chromatin modification to control flavonoid biosynthesis, providing a foundation for improving nutritional quality and stress resilience in banana.

## 1 Introduction

Plants synthesize a wide range of specialized metabolites that enable adaptation to dynamic environmental conditions and ecological interactions (Chen et al. 2022a, b). Among these, flavonoids constitute a diverse and evolutionarily conserved class of phenylpropanoid-derived compounds with essential roles in pigmentation, development, UV protection, and defense (Falcone Ferreyra et al. 2012; Naik et al. 2021). In addition to their physiological functions in plants, flavonoids confer significant health benefits due to their antioxidant, anti-inflammatory, and cardioprotective properties (Panche et al. 2016). Major subclasses, including anthocyanins and flavonols, contribute distinctly to plant fitness and stress adaptation (Shomali et al. 2022). Anthocyanins are responsible for red, purple, and blue pigmentation and function as antioxidants that protect against light-induced and oxidative stress while facilitating pollinator attraction (Mekapogu et al. 2020; Dong et al. 2023; Shen et al. 2024). Flavonols act as co-pigments, regulators of reactive oxygen species (ROS) homeostasis, and modulators of reproductive processes such as pollen germination (Zhang et al. 2009; Muhlemann et al. 2018). The flavonoid biosynthesis pathway is catalyzed by well-characterized enzymes, including CHS, CHI, F3H, F3′H, F3′5′H, FLS, and DFR representing a key regulatory step in anthocyanin production (Nagamatsu et al. 2007; Guo et al. 2019; Sun et al. 2021). At the transcriptional level, anthocyanin biosynthesis is primarily controlled by the MYB–bHLH–WD40 (MBW) complex, a conserved regulatory module across plant species (Rajput et al. 2022; Singh et al. 2024). Examples include PAP1/PAP2–EGL3/GL3–TTG1 in *Arabidopsis thaliana*, MdMYB10–MdbHLH3–MdTTG1 in apple, and MtLAP1–MtTT8–MtWD40-1 in *Medicago truncatula* (Zimmermann et al. 2004; Espley et al. 2007; Chagné et al. 2013; Li et al. 2016). In contrast, flavonol biosynthesis is often regulated independently of the MBW complex through subgroup 7 (SG7) R2R3-MYBs such as MYB11, MYB12, and MYB111, which directly activate early biosynthetic genes (Mehrtens et al. 2005; Stracke et al. 2007). However, emerging evidence suggests that additional regulators are required to fine-tune flavonoid metabolic flux in response to environmental cues.

B-box (BBX) transcription factors have recently emerged as key mediators linking light signaling to flavonoid biosynthesis (Li et al. 2025; Liang et al. 2025). These proteins, characterized by one or two zinc-finger B-box domains, are dynamically regulated by photoreceptors and environmental stimuli and bind G-box motifs in target promoters (Gangappa & Botto, 2014; Cao et al. 2023; Xiong et al. 2019; Song et al. 2020a). Functional studies demonstrate conserved roles for BBX proteins in promoting anthocyanin accumulation, as shown for IbBBX29 in sweet potato, VvBBX44 in grape, BBX16 in pear, and MdBBX37 in apple (Bai et al. 2019a; Liu et al. 2021; Gao et al. 2023). BBX proteins frequently interact with HY5 or MYB transcription factors to integrate environmental signals with pigment biosynthesis (Liu et al. 2022b; Fu et al. 2024). The BBX gene family plays a pivotal role in plant responses to abiotic stresses, including drought, cold, and heat stress (Li et al. 2025). For instance, SlBBX17 and SlBBX31 positively regulate thermotolerance in tomato, whereas AtBBX18 acts as a negative regulator of heat tolerance in *Arabidopsis* (Wang et al. 2013; Wang et al. 2024; Xu et al. 2022). The WRKY transcription factors constitute an additional regulatory layer linking specialized metabolism with stress-responsive signaling pathways. Depending on the biological context, these proteins can act either as activators or repressors. For instance, AtWRKY23 in *Arabidopsis thaliana* and NtWRKY11 in *Nicotiana tabacum* have been shown to positively regulate flavonoid biosynthesis (Grunewald et al. 2012; Wang et al. 2021). Similarly, MdWRKY11 and McWRKY71 function as positive regulators of anthocyanin and proanthocyanidin accumulation, whereas VvWRKY70 negatively regulates flavonol biosynthesis (Wang et al. 2018; Wei et al. 2023; Zhang et al. 2023). Despite these findings, how BBX and WRKY transcription factors integrate into coordinated regulatory networks governing flavonoid biosynthesis remains largely unresolved.

In addition to transcriptional control, epigenetic regulation plays a central role in modulating gene expression through dynamic changes in chromatin accessibility (Escrich et al. 2022). Environmental cues such as light and stress regulate histone acetyltransferases (HATs) and histone deacetylases (HDACs), leading to reversible chromatin remodeling (Perrella et al. 2016; Martínez-García & Moreno-Romero, 2020; Patitaki et al. 2022). Histone acetylation is generally associated with transcriptional activation, whereas deacetylation is linked to repression (Liu et al. 2022a; Chhatwal et al. 2024). For instance, UV-B exposure enhances histone acetylation in maize (Casati et al. 2008), while HDAC-mediated repression regulates anthocyanin biosynthesis under nutrient stress (Liao et al. 2022). Chromatin modifiers such as VvHDAC19 and HDA19 further regulate flavonoid-related genes through interactions with transcription factors (Jia et al. 2023; Jing et al. 2021), and chemical inhibition of HDACs enhances anthocyanin accumulation (Tang et al. 2025). In contrast, the role of histone acetyltransferases in flavonoid biosynthesis remains less defined. General Control Non-derepressible 5 (GCN5), a conserved GNAT-type HAT and core component of the SAGA complex, regulates plant development and stress responses through histone acetylation, including H3K9, H3K14, and H3K27 (Hu et al. 2015; Wang et al. 2016; Zheng et al. 2019; Kim et al. 2020; Tsilimigka et al. 2022). GCN5 also enhances tolerance to heat, salt, and drought stress by activating stress-responsive genes via chromatin remodeling (Hu et al. 2015; Li et al. 2019; Zheng et al. 2019). However, its direct role in flavonoid biosynthesis and its integration with transcriptional regulatory networks remain largely unexplored.

A major unresolved question is how light signaling coordinates transcriptional and epigenetic mechanisms to regulate tissue-specific flavonoid accumulation. Here, we identify a previously uncharacterized regulatory network centered on the B-box transcription factor MaBBX21 in banana. Functional analyses of overexpression and knockdown lines demonstrate that MaBBX21 positively regulates flavonoid biosynthesis. We further uncover a BBX–WRKY regulatory module in which MaBBX21 directly activates *MaWRKY23* to modulate flavonoid pathway flux. In addition, we reveal a novel role for the GNAT-type HAT MaGCN5 in specialized metabolism. *MaGCN5* acts downstream of MaBBX21 to promote H3K9 acetylation at the promoters of *MaDFR2, MaANS,* and *MaBBX21,* thereby reinforcing anthocyanin biosynthesis. Together, these findings define a MaBBX21–MaWRKY23/MaGCN5 regulatory axis that integrates light signaling with transcriptional and epigenetic control. This work provides a mechanistic framework for improving the nutritional quality and stress resilience of banana.

## 2 Results

### 2.1 Transcriptome Analysis of Bract, Flower, and Leaf Tissue of Banana Unveiled Distinct Transcriptional Programming Governing Flavonoids Accumulation

Banana tissues exhibit distinct flavonoid profiles, with anthocyanins predominantly accumulating in bracts, while flowers contain higher total phenolics but lack visible pigmentation (Chiang et al. 2021). Leaves, in contrast, show the highest flavonol accumulation (Pandey et al. 2016). Consistently, LC–MS analysis revealed elevated levels of catechin, procyanidin B1/C1, quercetin, rutin, and kaempferol-3-O-rutinoside in leaves, whereas cyanidin was the major anthocyanidin in bracts (Figure 1a, S1a–c), with significantly higher total anthocyanin levels in bracts (Figure 1b). This divergence suggests differential pathway regulation. Expression profiling supported this, showing that many early biosynthesis genes were enriched in leaves and flowers, while late biosynthesis genes (*MaDFR2*, *MaANS*) were strongly and specifically expressed in bracts, correlating with anthocyanin accumulation (Figure S2a, b). To get insight into the transcriptional landscape behind this spatial gene expression profile of FBGs, we performed transcriptome analysis of these tissues. RNA-seq analysis further revealed clear tissue-specific patterns, with PCA distinctly separating leaves from bract and flower tissues (Figure 1c). Differential expression analysis (|log FC| ≥ 1; adjusted p < 0.05) identified substantial numbers of DEGs across comparisons (Figure S3a), with overlaps illustrated in Venn diagrams (Figure S3b). GO and KEGG analyses indicated that bract-enriched genes were associated with flavonoid biosynthesis and light/hormone responses, leaf-enriched genes with photosynthesis and stress pathways, and flower-enriched genes with carbohydrate metabolism and stress responses (Figure S4, S5). Regulatory gene expression also showed tissue specificity. Among these, BBX family is widely implicated to play critical role in light signalling pathway to modulate various metabolic and developmental pathway. Several *MaBBX* genes were differentially enriched in leaves and bracts (Figure 1d, S6a), while *MaHY5* expression was higher in leaves (Figure 1e). Apart from these, photoreceptors such as *MaPIF8* were expressed in both bracts and leaves, whereas *MaCRY1/2* were more abundant in bracts and flowers (Figure S6b). Additionally, *MaWRKY* genes displayed antagonistic expression patterns between leaf and bract tissues (Figure S6c), suggesting their role in tissue-specific flavonoid regulation.

**FIGURE 1.**
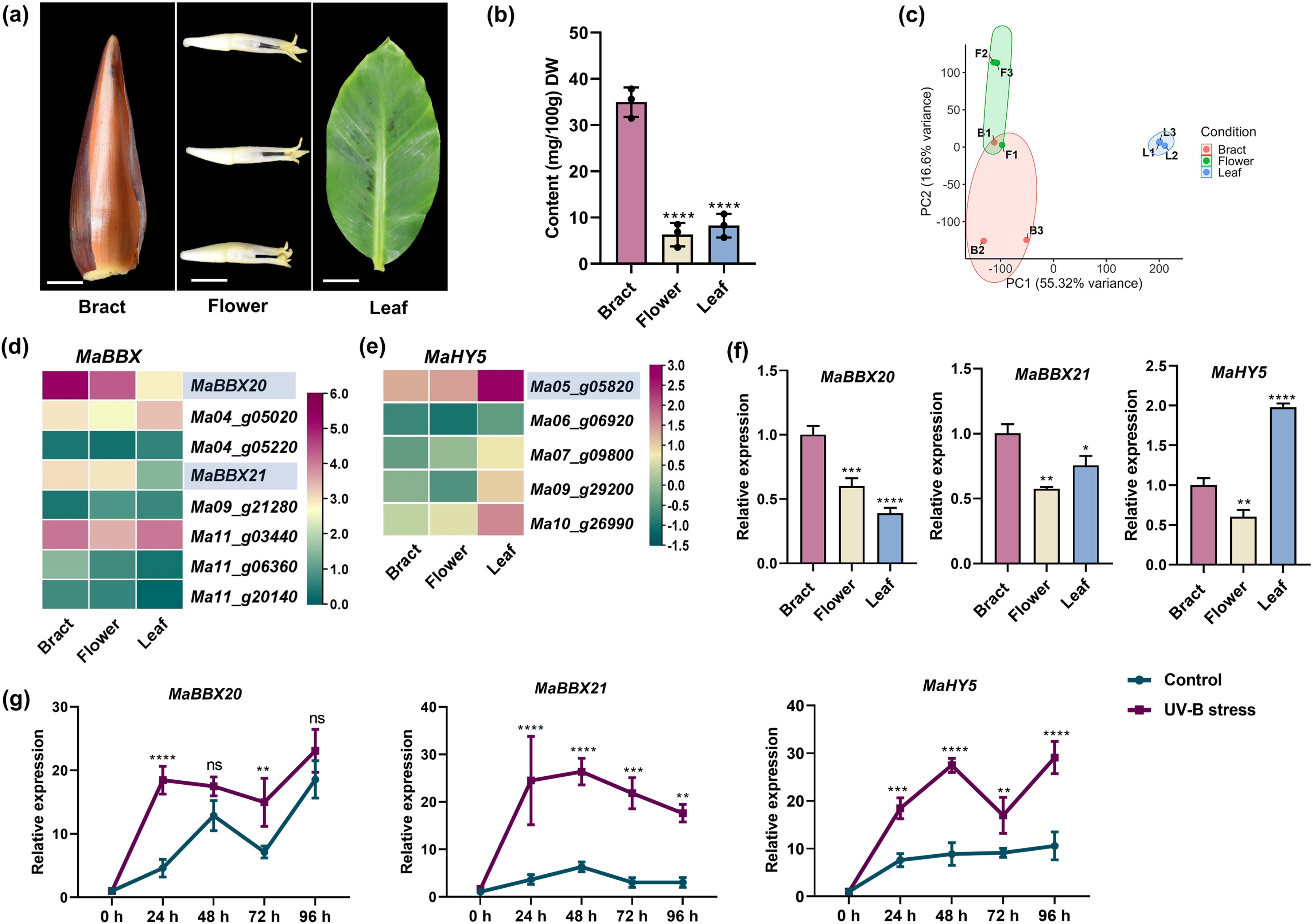
Transcriptome analysis of bract, flower and leaf tissues and expression profiling of *MaBBX* and *MaHY5* genes. (a) Pictorial representation of bract, flower, and leaf tissues highlighting visible phenotypic differences in pigmentation. (b) The bar graph illustrates the total anthocyanin content across these tissues, showing significant variation. (c) Principal Component Analysis (PCA) plot illustrating distinct gene expression profiles in these tissues. The separation along PC1 (55.32% variance) highlights significant transcriptional variation, particularly distinguishing leaf tissue from bract and flower. (d) Heatmaps illustrate the normalized expression patterns of selected *MaBBX* (members of subgroup-IV). (e) Heatmaps showing the normalized expression patterns of *MaHY5* in the transcriptome dataset. (f) RT–qPCR-based expression analysis of *MaBBX20, MaBBX21,* and *MaHY5* across bract, flower, and leaf tissues. (g) Relative expression profiling of *MaBBX20, MaBBX21,* and *MaHY5* in leaf tissue of banana saplings at 0 h, 24 h, 48 h, 72 h and 96 h under control and UV-B stress conditions, as determined by RT–qPCR, showing differential transcriptional responses. Asterisks represent statistically significant differences (\**p* < 0.05, \*\**p* < 0.01, \*\*\**p* < 0.001, \*\*\*\**p* < 0.0001, ns = not significant).

### 2.2 *MaBBX* Genes Might be Playing Critical Roles in the Anthocyanin Accumulation in the Different Colour Contrasting Tissues

Flavonol and anthocyanin accumulation is prominent in light-exposed tissues such as bracts and leaves, suggesting involvement of light signaling pathways. Multiple transcription factors regulate light-induced flavonoid biosynthesis (Naik et al. 2022). Due to the involvement of BBX family genes in light induced flavonoids biosynthetic pathway and their greater transcript enrichment in bract tissue, we focussed on members of this gene family. Phylogenetic analysis of *MaBBX* proteins, using characterized BBXs from *Arabidopsis*, rice, and maize, classified them into five clades (I–V) (Figure S7a). A subset within clade IV, associated with flavonoid regulation, comprised eight members with conserved two B-box domains (Figure S7b), but displayed distinct tissue-specific expression patterns (Figure 1d, f; Figure S7c), indicating functional diversification.

Among these, *MaBBX20* (Ma01_g03030) and *MaBBX21* (Ma07_g18870) showed higher expression in anthocyanin-rich bracts, moderate levels in flowers, and low expression in leaves, as confirmed by RT-qPCR (Figure 1f), suggesting their role in anthocyanin accumulation. Given that BBXs respond to UV-B and regulate anthocyanin biosynthesis (Podolec et al. 2020; Job et al. 2022), expression analysis under UV-B revealed strong and sustained induction of *MaBBX21* and *MaHY5*, whereas *MaBBX20* showed weaker responsiveness (Figure 1g). Subcellular localization confirmed nuclear localization of both proteins (Figure S7d). Transient overexpression assays further showed that both *MaBBX20* and *MaBBX21* activate anthocyanin biosynthetic genes (*MaDFR2*, *MaANS*), with *MaBBX21* exhibiting a stronger effect (Figure S8, S9). Together, these findings identify *MaBBX21* as a key light-responsive regulator of flavonoid biosynthesis, warranting detailed functional characterization.

### 2.3 *MaBBX21* Promotes Flavonol and Anthocyanin Biosynthesis in Banana

To assess the role of *MaBBX21*, transient overexpression (OE) and knockdown (KD) were performed in banana fruit discs. Transformation efficiency was confirmed by GUS staining and RT-qPCR, showing strong induction (∼491-fold) in OE and suppression (∼0.4-fold) in KD tissues (Figure S9a, b). OE significantly upregulated several flavonoid biosynthetic genes (*MaCHS*, *MaCHI*, *MaF3H*, *MaFLS*, *MaF3*′*5*′*H*), while KD substantially repressed them. Notably, anthocyanin-specific genes (*MaDFR2*, *MaANS*) were strongly induced in OE and reduced in KD (Figure S9b). These changes correlated with metabolite accumulation, where flavonols (kaempferol, myricetin) and anthocyanins (cyanidin, delphinidin, petunidin, peonidin) increased in OE and decreased in KD (Figure S9c).

Stable *MaBBX21*-OE and KD lines were generated and validated by RT-qPCR and GUS staining (Figure S10a–d), with selected lines showing maximum change in expression levels (Figure 2a). OE plants exhibited red pigmentation in leaves and pseudostems, whereas KD lines lacked pigmentation (Figure 2b, c). Expression of key biosynthetic genes (*MaFLS1/2*, *MaF3’H*, *MaDFR1/2*, *MaANS*) was upregulated in OE and downregulated in KD lines (Figure 2d; Figure S11a, b). Consistently, DPBA staining showed enhanced flavonol accumulation in OE roots and reduced signals in the roots of KD lines (Figure 2e). Metabolite analysis further confirmed these trends, with total flavonols (sum of quercetin, kaempferol, myricetin) and anthocyanins increasing by ∼4-6-fold and ∼6-8-fold, respectively, in OE lines and decreasing in KD plants (Figure 2f). LC–MS profiling supported increased accumulation of major flavonols (quercetin, kaempferol, myricetin, rutin) and anthocyanidins (cyanidin, delphinidin, petunidin, peonidin, pelargonidin) in OE lines, with opposite trends in KD as compared to control (Figure S12a, b). Collectively, these results establish *MaBBX21* as a central positive regulator of flavonoid biosynthesis in banana.

**FIGURE 2.**
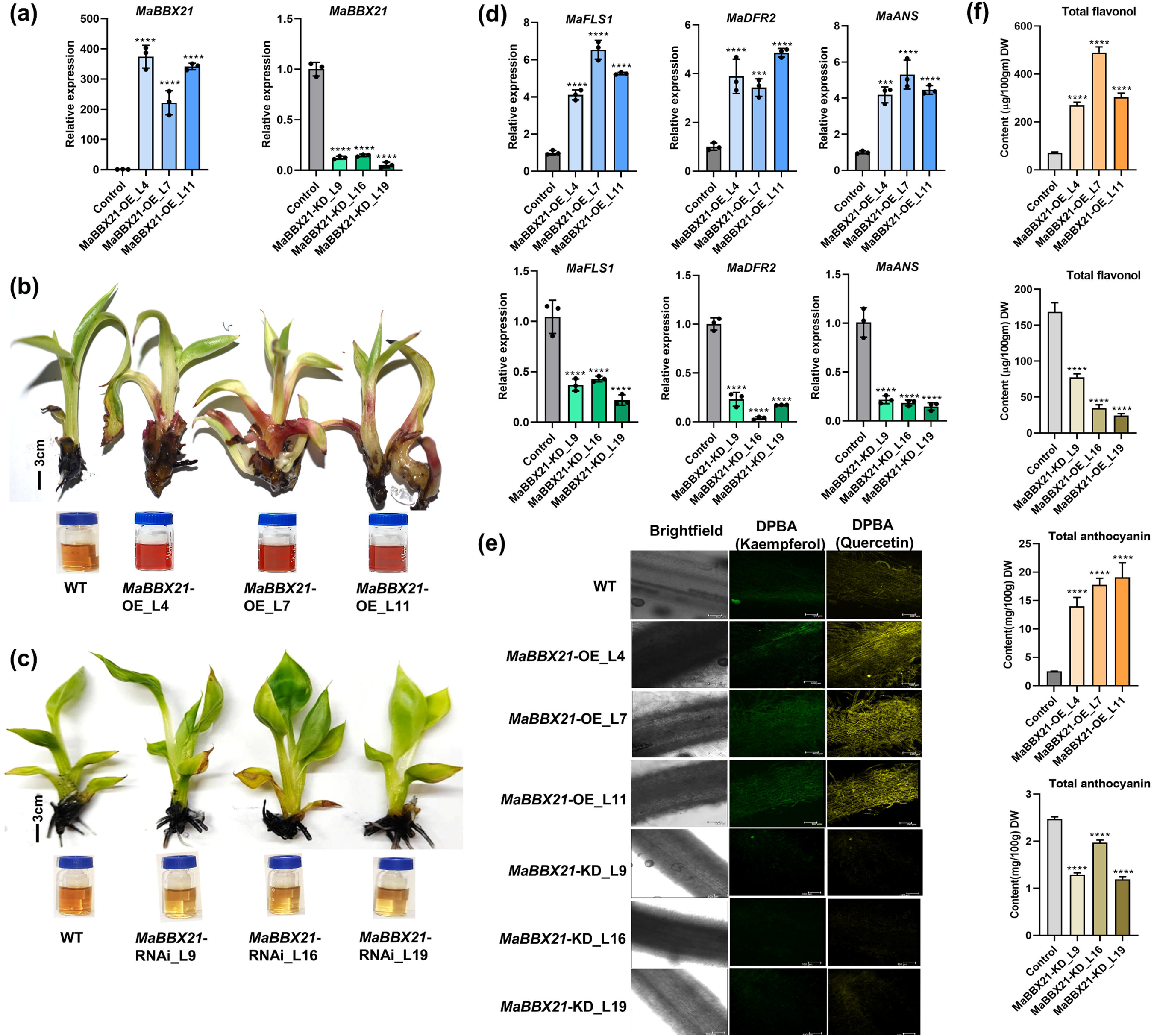
*MaBBX21* positively regulates flavonol and anthocyanin accumulation in banana plants. (a) Relative expression levels of *MaBBX21* in three independent OE and KD banana lines, confirmed by RT–qPCR. Expression levels were normalized to the internal control and compared with WT plants. (b) Representative phenotypes of WT and three independent *MaBBX21*-OE lines showing enhanced red pigmentation and visibly increased total anthocyanin accumulation compared with WT. (c) In contrast, WT and three *MaBBX21*-KD lines displayed no obvious visible pigmentation differences but visibly reduced total anthocyanin content in KD lines. (d) Relative transcript levels of key flavonoid biosynthesis genes *(MaFLS1 MaDFR2,* and *MaANS*) in WT, *MaBBX21*-OE and KD lines as determined by RT–qPCR. Expression levels were calculated relative to the corresponding WT plants. (e) DPBA staining of banana roots showing flavonol accumulation in *MaBBX21*-OE lines which exhibit stronger green fluorescence (kaempferol) and yellow fluorescence (quercetin) relative to WT, whereas KD lines show reduced fluorescence intensity. (f) Quantification of total flavonol content (sum of quercetin, myricetin, and kaempferol) and total anthocyanin levels in *MaBBX21*-OE and *MaBBX21*-KD banana lines. Total anthocyanin is expressed as mg, while total flavonol content is presented as µg per 100 g dry weight (DW). Data represent the mean ± SD of three independent biological lines, with three technical replicates. Statistical significance was determined using one-way ANOVA followed by Dunnett’s multiple comparisons test Asterisks indicate significance levels: \**p* ≤ 0.05, \*\**p* ≤ 0.01, \*\*\**p* ≤ 0.001, and \*\*\*\**p* ≤ 0.0001; ns, not significant.

### 2.4 MaBBX21 Cooperates with MaHY5 to Activate Anthocyanin Biosynthesis Genes

BBX–HY5 interactions are conserved regulators of light-responsive gene expression (Job et al. 2022). Consistently, protein–protein interaction assays (Y2H, BiFC, and LCI) confirmed a strong and specific interaction between MaBBX21 and MaHY5, whereas the homolog MaBBX20 showed no interaction with MaHY5 (Figure 3a-c; Figure S13a, b), indicating functional specificity.

**FIGURE 3.**
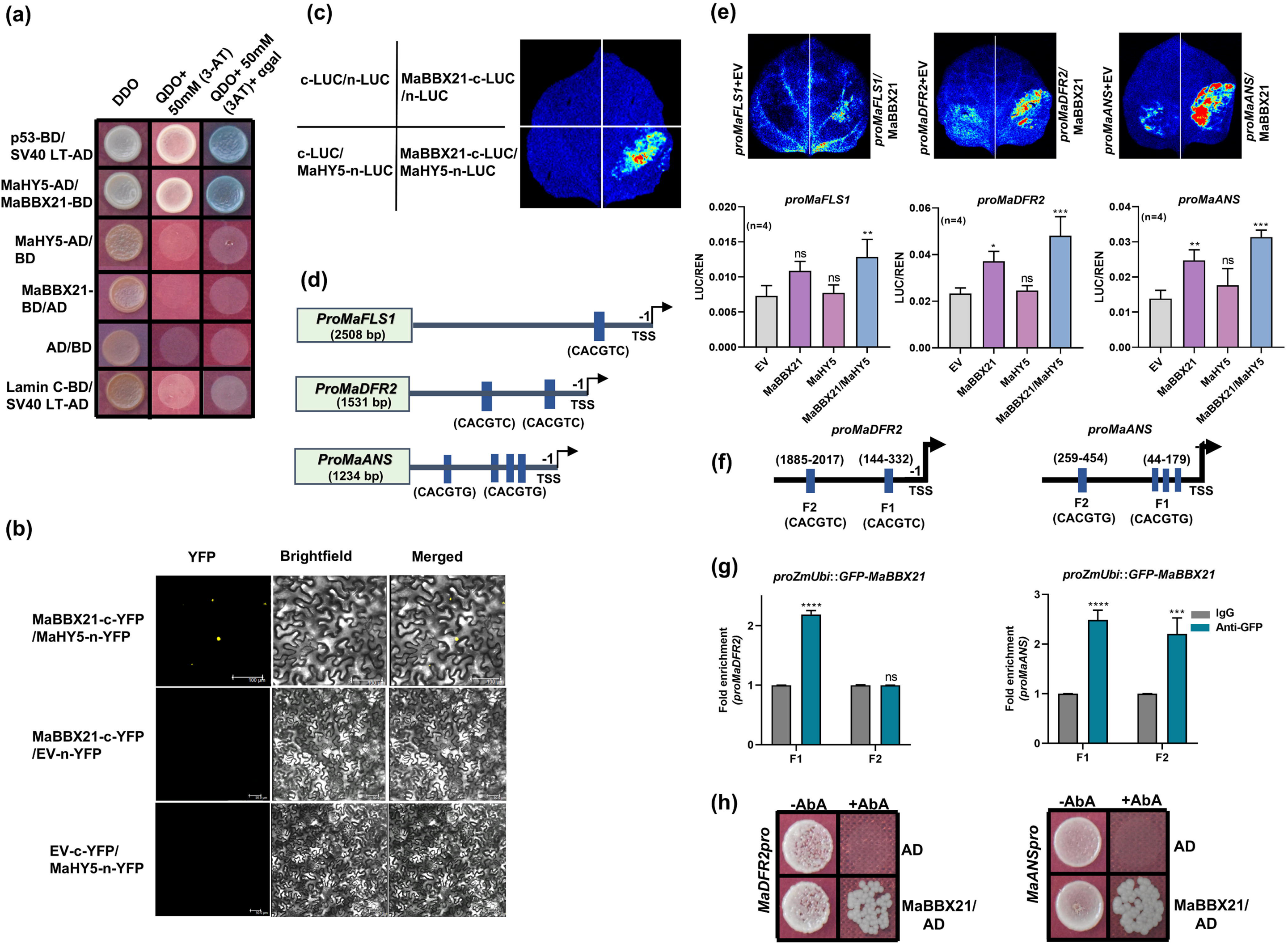
MaBBX21 interacts with MaHY5 to activate *MaDFR2* and *MaANS* for anthocyanin biosynthesis. (a) Yeast two-hybrid (Y2H) assay showing that MaBBX21 specifically interacts with MaHY5. MaBBX21 was fused to the GAL4 activation domain (BD), while MaHY5 were fused to the GAL4 DNA-binding domain (AD). Yeast transformants were selected on double dropout (DDO; -Leu/-Trp) medium and screened for interaction on quadruple dropout (QDO; - Leu/-Trp/-Ade/-His) medium. Strong interactions were confirmed by the formation of blue colonies on QDO medium supplemented with X-α-Gal. (QDO/X/3-AT), due to α-glucosidase activity. (b) Bimolecular fluorescence complementation (BiFC) assay confirming *in-*planta interaction between MaBBX21 and MaHY5 in *N. benthamiana* leaves. Fluorescence signal indicates protein-protein interaction. YFPn + YFPc was used as a negative control. Scale bar = 20 µm. (c) Luciferase complementation imaging (LCI) assay showing luminescence signals resulting from the interaction of MaBBX21 with MaHY5 (MaBBX21-cLUC + MaHY5-nLUC) *in*-planta. Negative controls included MaBBX21-cLUC + nLUC, cLUC + MaHY5-nLUC, and nLUC + cLUC, which showed no luminescence. (d) Schematic diagram of the promoter regions of *MaFLS1, MaDFR2,* and *MaANS,* highlighting putative BBX binding motifs. (e) Transactivation analysis of *proMaFLS1–2508, proMaDFR2-1531*, and *proMaANS-1234,* promoters by MaBBX21, MaHY5, using dual-luciferase assays in *N. benthamiana* leaves. Results are shown as LUC/REN ratios, representing firefly luciferase normalized to Renilla luciferase activity. Data are presented as mean ± SD from four independent biological replicates (n = 4). (f) Promoter fragments of *MaDFR2* and *MaANS* upstream of the transcription start site (TSS) are shown, with predicted BBX binding motifs indicated by blue boxes. The positions of the ChIP–qPCR primer–amplified regions are also indicated to assess MaBBX21 enrichment at these promoters. (g) ChIP-qPCR was performed to assess the *in vivo* binding of MaBBX21 to the promoters of *MaDFR2,* and *MaANS,* promoters. Fold enrichment is shown as mean ± SD from two biological replicates. Standard deviation and significance level are represented by error bars and asterisks, respectively. Asterisks represent statistically significant differences (\**p* < 0.05, \*\**p* < 0.01, \*\*\**p* < 0.001, \*\*\*\**p* < 0.0001, ns = not significant). (h) Yeast one hybrid assay demonstrating the binding of MaBBX21 to promoter fragments of*, MaDFR2,* and *MaANS* containing BBX binding motifs. An empty pGADT7 vector (EV) was used as a negative control. Successful protein-DNA interaction was indicated by yeast growth on SD/−Ura media supplemented with aureobasidin A (AbA). AD = activation domain.

Promoter analysis of *MaFLS1*, *MaDFR2*, and *MaANS* identified conserved BBX-binding motifs (CACGTC/G) (Figure 3d). Dual-luciferase assays revealed that *MaBBX21* directly activates *MaDFR2* and *MaANS*, with enhanced activation in the presence of *MaHY5*, while *MaHY5* alone showed minimal activity. In contrast, *MaFLS1* activation required co-expression of both factors, indicating cooperative regulation (Figure 3e). ChIP–qPCR using ZmUBIpro::*MaBBX21*-GFP confirmed *in vivo* enrichment at *MaDFR2* and *MaANS* promoters (Figure 3f-g), further supported by Y1H assays, demonstrating direct binding (Figure 3h). Together, these results establish MaBBX21 as a direct activator of anthocyanin biosynthesis genes, with MaHY5 acting as a co-regulator, while flavonol regulation likely occurs indirectly within the BBX-mediated network.

### 2.5 *MaWRKY23* Functions as a Downstream Target of MaBBX21 to Enhance Flavonol Accumulation

Given that flavonols also accumulate abundantly in leaves (Pandey et al. 2016), and that DFR and FLS compete for dihydroflavonol to direct flux toward anthocyanin or flavonol branches, *MaBBX21* likely modulates pathway flux in a tissue-dependent manner. To identify downstream regulators, expression of key flavonoid-related transcription factors, including MYB, bHLH, WD40, and WRKY families, was analyzed in *MaBBX21*-OE and KD lines. Among these, *MaWRKY23* showed a strong and specific response, with 4-6-fold upregulation in OE lines and significant repression (0.2–0.4-fold) in KD lines (Figure 4a; Figure S14a, b). In contrast, most *MaMYB*, *MabHLH*, *MaWD40*, and other *MaWRKY* genes showed minimal changes, although *MaMYBPA2* was downregulated in KD lines (Figure S14a, b). These results indicate that *MaBBX21* primarily regulates flavonol biosynthesis via *MaWRKY23* instead of conventional MBW complex regulation.

**FIGURE 4.**
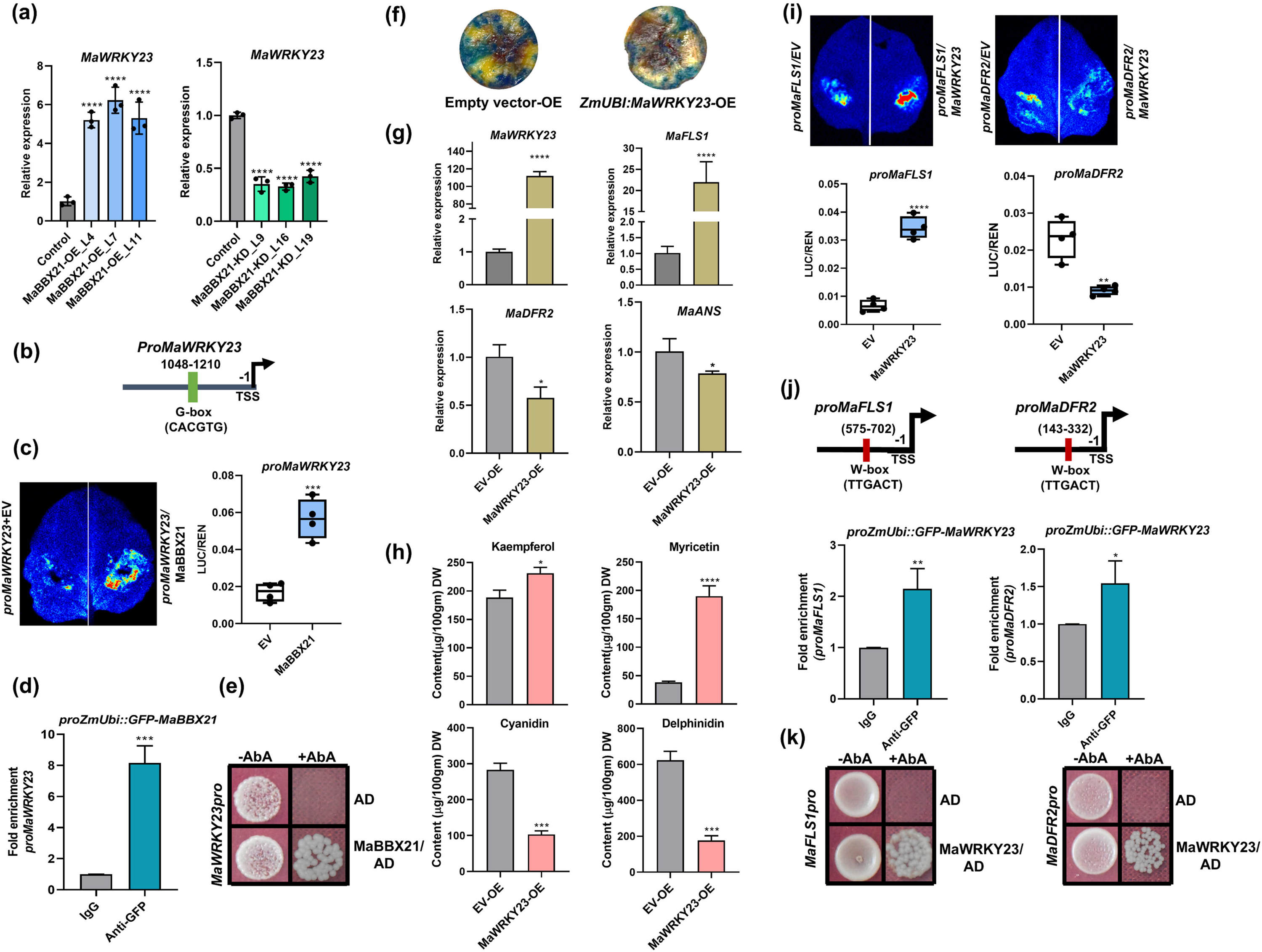
*MaWRKY23* functions as a downstream target of MaBBX21 to enhance flavonol accumulation. (a) Relative expression of *MaWRKY23* in *MaBBX21*-OE and KD banana lines. (b) Promoter analysis of *MaWRKY23* reveals the presence of a single putative MaBBX21-binding site upstream of the transcription start site. (c) Dual-luciferase reporter assays demonstrate that MaBBX21 positively transactivates the *MaWRKY23* promoter, indicating direct transcriptional regulation. (d) ChIP-qPCR was performed to assess *in-vivo* binding of MaBBX21 to the promoter of *MaWRKY23*. (e) Yeast one hybrid assay demonstrating the binding of MaBBX21 to promoter fragments of *MaWRKY23* containing BBX binding motif. (f) Representative images of GUS-stained banana fruit discs confirm successful transient transformation with *MaWRKY23* OE. Images were taken 3 d post-transformation. (g) The relative expression of *MaWRKY23*, *MaFLS1, MaDFR2* and *MaANS* was analysed in transiently *MaWRKY23-*OE banana fruit discs using RT-qPCR. Expression levels were compared to EV control. (h) The accumulation of flavonols (kaempferol and myricetin), and anthocyanin derivatives (cyanidin and delphinidin) was quantified in *MaWRKY23*-OE banana fruit discs. (i) Transactivation assay of *proMaFLS1–*2508 and *proMaDFR2-*1531 by MaWRKY23 using dual-luciferase assays in *N. benthamiana* leaves; n=4. (j) The positions of W-box (red line) and the ChIP–qPCR primer–amplified regions are shown in the promoters of *MaFLS1* and *MaDFR2*. ChIP-qPCR was performed to assess the *in-vivo* binding of MaWRKY23 to W-box element within these promoters. Standard deviation and significance level are represented by error bars and asterisks, respectively. Asterisks represent statistically significant differences (\**p* < 0.05, \*\**p* < 0.01, \*\*\**p* < 0.001, \*\*\*\**p* < 0.0001, ns = not significant). (k) Yeast one hybrid assay demonstrating the binding of MaWRKY23 to promoter fragments of *MaFLS1* and *MaDFR2,* containing W-box binding motifs. Successful protein-DNA interaction was indicated by yeast growth on SD/−Ura media supplemented with aureobasidin A (AbA). AD = activation domain.

Sequence analysis of the *proMaWRKY23* (1.7 kb) identified a conserved BBX-binding motif (CACGTG) ∼1152–1158 bp upstream of the TSS, suggesting direct regulation by MaBBX21 (Figure 4b). Dual-luciferase assays confirmed strong activation of *proMaWRKY23* by MaBBX21 (Figure 4c), which was further validated by *in vivo* ChIP–qPCR and Y1H assays demonstrating direct binding (Figure 4d, e). Functional characterization showed that transient OE of *MaWRKY23* in banana fruit discs led to ∼112-fold induction of its transcript. This resulted in differential regulation of flavonoid biosynthesis genes, with upregulation of *MaCHS5/6*, *MaCHI2*, *MaF3H2*, and *MaF3*′*5*′*H4/6*, while *MaCHS1/2*, *MaCHI1*, *MaF3*′*5*′*H1–3*, *MaDFR2*, and *MaANS* were repressed. Notably, *MaFLS1* was strongly induced (∼21-fold), accompanied by increased flavonol accumulation (myricetin, kaempferol) and reduced anthocyanin levels (Figure 4f–h; Figure S15), indicating a metabolic shift toward flavonol biosynthesis. Cis-element analysis of *MaFLS1* and *MaDFR2* promoters revealed W-box motifs (TTGACG), typical WRKY binding sites. Dual-luciferase, ChIP–qPCR, and Y1H assays confirmed that *MaWRKY23* directly binds these promoters, activating *proMaFLS1* while repressing *proMaDFR2* (Figure 4i–k). Collectively, these results establish *MaWRKY23* as a crucial regulator that redirects metabolic flux toward flavonol biosynthesis, functioning downstream of *MaBBX21*.

### 2.6 *MaGCN5* Acts Downstream of MaBBX21 to Enhance H3K9 Acetylation

The spatiotemporal pattern of anthocyanin accumulation in *MaBBX21*-OE plants suggested involvement of epigenetic regulation alongside transcriptional control. Given that light enhances histone acetylation, particularly H3K9ac (Perrella et al. 2016), we hypothesized that *MaBBX21* regulates anthocyanin biosynthesis via chromatin modification. Western blot analysis revealed significantly elevated global H3K9ac levels in *MaBBX21*-OE plants compared to WT (Figure 5a). Consistent with reports linking H3K9ac to flavonoid gene activation (Li et al. 2020; Patrick et al. 2021), ChIP–qPCR showed strong enrichment of H3K9ac at the *proMaDFR2* and *proMaANS* regions (P1/P2) in *MaBBX21*-OE lines (Figure 5b), indicating locus-specific activation. Out of four types of histone acetyltransferases (HAT) classes known so far, GNAT-type are thoroughly studied in plants, particularly under stress. Among GNAT-type histone acetyltransferases, *MaGCN5* exhibited a strong response, with 6–9-fold upregulation in *MaBBX21*-OE lines and repression in KD plants, while *HAG2* and *HAG3* showed minimal changes (Figure 5c; Figure S16a). Promoter analysis identified BBX-binding motifs in *proMaGCN5*, and dual-luciferase assays confirmed its activation by MaBBX21 (Figure 5d). This was supported by ChIP–qPCR and Y1H assays demonstrating direct binding of *MaBBX21* to the F1 region of *proMaGCN5* (Figure 5e, f). Subcellular localization further confirmed nuclear localization of *MaGCN5* (Figure S16b). Pharmacological inhibition using γ-butyrolactone reduced *MaGCN5* expression and suppressed *MaDFR2* and *MaANS* transcription even in *MaBBX21*-OE lines (Figure 5g), confirming its functional role. Collectively, these results demonstrate that MaBBX21 directly activates *MaGCN5*, which enhances H3K9 acetylation at flavonoid biosynthesis gene promoters, thereby promoting their expression.

**FIGURE 5.**
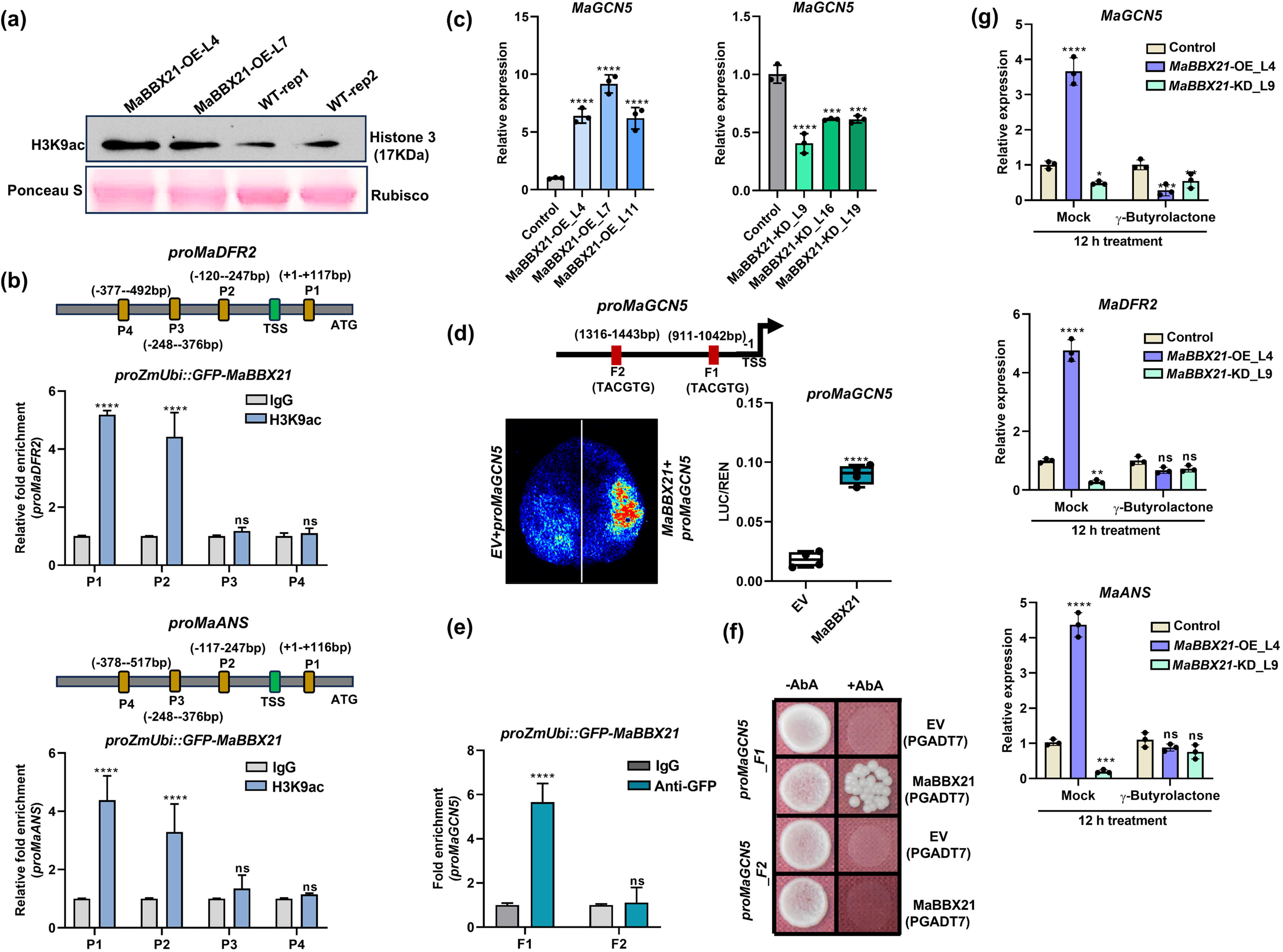
MaBBX21 enhances *MaGCN5* expression by binding to its promoter and enhances acetylation level. (a) Western blot analysis using an anti-H3K9ac antibody revealed elevated global H3K9 acetylation levels in *MaBBX21*-OE plants compared with WT plants. Ponceau S staining serves as a control for equal protein loading and transfer efficiency. (b) ChIP–qPCR analysis with an H3K9ac-specific antibody confirmed enhanced enrichment of H3K9 acetylation at the promoters of *MaDFR2* and *MaANS* in *MaBBX21*-OE plants. Four fragments (P1 to P4) near TSS are shown with mustard yellow boxes on *proMaDFR2* and *proMaANS*. (c) RT–qPCR analysis showed that the histone acetyltransferase *MaGCN5* was significantly upregulated in *MaBBX21*-OE lines and downregulated in *MaBBX21*-KD plants. (d) Two putative BBX-binding *cis-*elements are marked with red boxes within the *MaGCN5* promoter, and dual-luciferase assays demonstrated that MaBBX21 positively transactivates the *MaGCN5* promoter. (e) ChIP–qPCR revealed significant enrichment of MaBBX21 at the F1 fragment of *proMaGCN5*, whereas no enrichment was detected at the F2 fragment. (f) Yeast one-hybrid assays further validated the direct interaction between MaBBX21 and the F1 fragment of *proMaGCN5*. (g) Chemical inhibition of MaGCN5 activity in *MaBBX21*-OE, KD, and WT plants significantly reduced *MaGCN5* transcript levels. Consequently, expression of the flavonoid biosynthesis genes *MaDFR2* and *MaANS* was markedly decreased upon inhibitor treatment. Standard deviation and significance level are represented by error bars and asterisks, respectively. Asterisks represent statistically significant differences (\**p* < 0.05, \*\**p* < 0.01, \*\*\**p* < 0.001, \*\*\*\**p* < 0.0001, ns = not significant).

### 2.7 MaGCN5 Occupies and Acetylates H3K9 At the Promoter Regions of *MaDFR2*, *MaANS* and *MaBBX21* to Promote Anthocyanin Accumulation

Elevated expression of flavonoid biosynthesis genes together with increased H3K9ac at their TSS-proximal regions in *MaBBX21*-OE lines suggested a regulatory role for *MaGCN5* in flavonoid biosynthesis. To validate this, *MaGCN5* was transiently OE and KD in banana fruit discs and embryogenic cell suspension (ECS) and confirmed by GUS staining (Figure 6a, b). *MaGCN5*-OE resulted in ∼13-fold (fruit discs) and ∼9-fold (ECS) transcript induction, whereas KD reduced expression to ∼0.3-fold and ∼0.1-fold, respectively. Functional analyses showed that *MaGCN5*-OE significantly upregulated key structural genes, including *MaDFR2*, *MaANS*, and *MaFLS1*, while these genes were repressed in KD samples (Figure 6c, d). In contrast, *MaWRKY23*, *MaFLS2*, and *MaF3*′*H* displayed inconsistent expression patterns in fruit (Figure S17a) and callus (Figure S17b). Metabolite profiling corroborated these results, with increased anthocyanin accumulation (cyanidin, peonidin, pelargonidin) in OE tissues and reduced levels in KD samples (Figure 6e, f).

**FIGURE 6.**
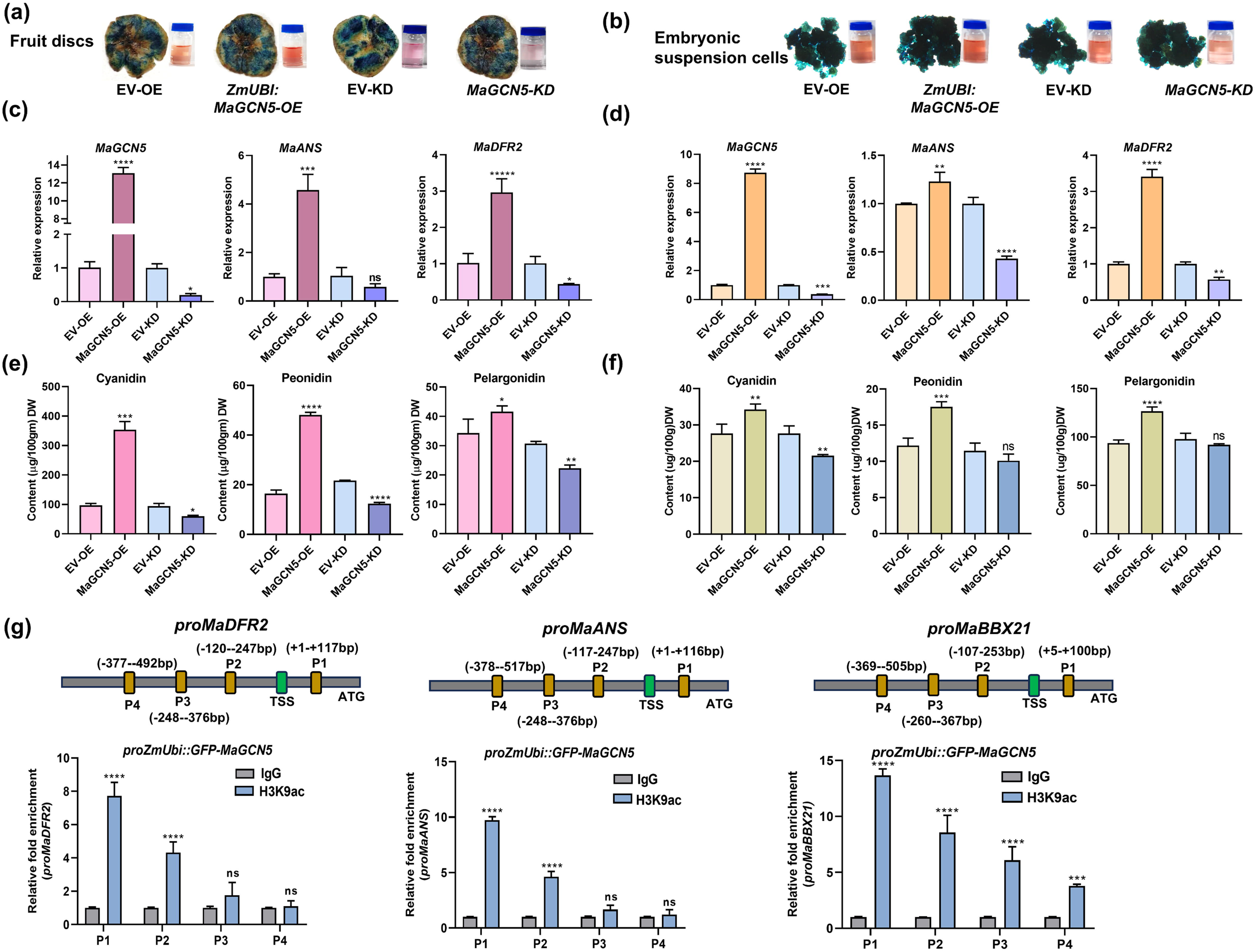
MaGCN5 enhances anthocyanin accumulation through H3K9 acetylation at *MaANS* and *MaDFR2* promoters. (a) Representative image of GUS-stained banana fruit discs and (b) Embryonic cells suspension (ECS) showing successful transient OE and KD of *MaGCN5*. Visual differences in anthocyanin accumulation are evident among OE, KD and their respective controls. (c) Relative transcript levels of *MaGCN5* and flavonoid biosynthesis genes (*MaDFR2* and *MaANS*) in *MaGCN5*-OE and KD banana fruit discs and (d) ECS, determined by RT–qPCR analysis. (e) Quantification of anthocyanidin derivatives (cyanidin, peonidin and pelargonidin) in *MaGCN5*-OE and KD banana fruit discs and (f) ECS, determined by LCMS. (g) Four promoter fragments (P1–P4) located near the TSS are highlighted by mustard yellow boxes in *proMaDFR2*, *proMaANS* and *proMaBBX21*. ChIP–qPCR analysis of histone H3K9 acetylation levels at the *MaDFR2*, *MaANS* and *MaBBX21* promoter loci using an anti-H3K9ac antibody in *MaGCN5*-OE ECS samples. Standard deviation and significance level are represented by error bars and asterisks, respectively. Asterisks represent statistically significant differences (\**p* < 0.05, \*\**p* < 0.01, \*\*\**p* < 0.001, \*\*\*\**p* < 0.0001, ns = not significant).

ChIP–qPCR using GFP–*MaGCN5* demonstrated significant enrichment at TSS-proximal (P1/P2) regions of *MaDFR2*, *MaANS*, and *MaBBX21* promoters, indicating direct occupancy at active loci (Figure S18). Correspondingly, H3K9ac ChIP–qPCR revealed strong enrichment at *proMaDFR2* and *proMaANS*, confirming that *MaGCN5* catalyzes histone acetylation to activate transcription. Notably, enhanced H3K9ac was also observed in all the (P1-P4) *MaBBX21* promoter (Figure 6g), indicating epigenetic activation of *MaBBX21* itself. Together, these findings support a positive feedback loop in which *MaBBX21* activates *MaGCN5*, and *MaGCN5* in turn enhances *MaBBX21* expression via H3K9 acetylation, collectively amplifying flavonoid biosynthesis in banana.

### 2.8 *MaBBX21* Increases Heat Tolerance by Stimulating Flavonoid Biosynthesis and Reducing ROS Accumulation

BBX proteins play crucial roles in signaling cascades under diverse abiotic stresses. To investigate the function of *MaBBX21* in heat stress, two-month-old WT, *MaBBX21*-OE, and KD plants were subjected to elevated temperature conditions. After 72 h of heat treatment, *MaBBX21*-OE lines exhibited enhanced thermotolerance, as evidenced by reduced tissue damage and fewer burn lesions, whereas KD plants displayed burn symptoms comparable to WT (Figure 7a). At the transcriptional level, heat stress induced the expression of *MaBBX21, MaWRKY23, MaGCN5,* and *MaFLS1* in WT plants; however, their expression was reduced in KD lines. In contrast, *MaDFR2* and *MaANS* were generally downregulated in both OE and KD plants (Figure 7b; Figure S19a).

**FIGURE 7.**
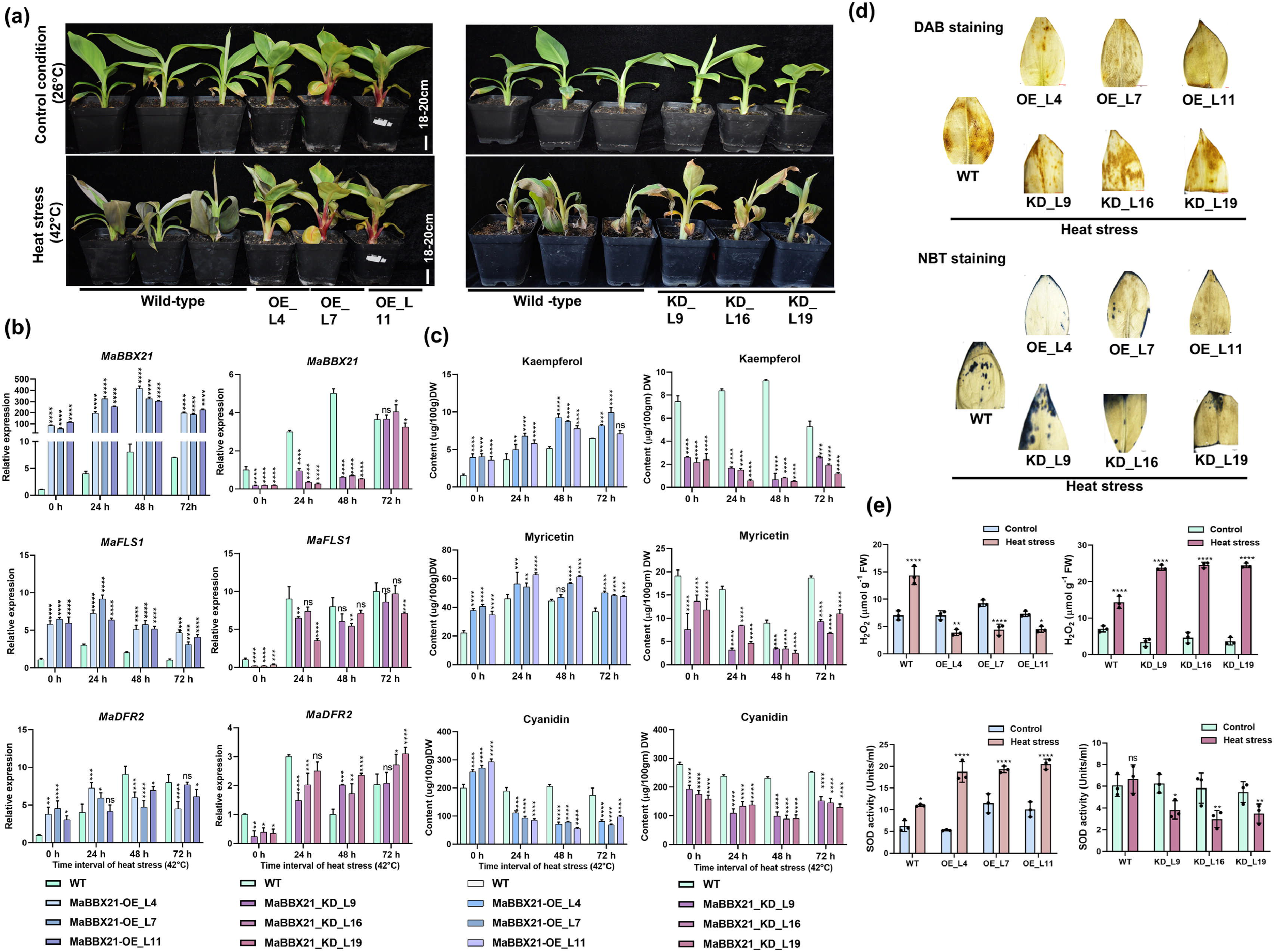
*MaBBX21* enhances heat tolerance by promoting flavonoid biosynthesis and restricting ROS accumulation. (a) Phenotypic comparison of two-month-old WT and *MaBBX21*-OE and KD saplings grown at 26 °C and subsequently exposed to heat stress (42 °C) for 72 h. (b) Expression analysis of *MaBBX21, MaFLS1* and *MaDFR2*, and (c) quantification of flavonoid content in WT, *MaBBX21*-OE and KD banana lines at 0 h, 24 h, 48 h, and 72 h of heat stress. Data represent mean ± SD of WT and three independent lines, each with three technical replicates. (d) DAB and NBT staining in WT, *MaBBX21*-OE and KD lines after 48 h of heat stress. (e) Estimation of H O content and SOD enzyme activity in WT, *MaBBX21*-OE and KD lines at 48 h of heat stress. Asterisks represent statistically significant differences (\**p* < 0.05, \*\**p* < 0.01, \*\*\**p* < 0.001, \*\*\*\**p* < 0.0001, ns = not significant).

Metabolite profiling revealed a stress-induced shift toward flavonol accumulation. In WT plants, kaempferol, myricetin, and rutin levels increased across all time points, with further enhancement observed in OE lines and a reduction in KD lines. Quercetin accumulation increased at early time points but declined at later stages in OE plants, while it remained consistently lower in KD lines compared to WT. Conversely, anthocyanin derivatives, including cyanidin and delphinidin, were reduced in both OE and KD lines, indicating a reprogramming of metabolic flux (Figure 7c; Figure S19b). Enhanced thermotolerance in OE lines was associated with improved redox homeostasis. Histochemical staining (DAB and NBT) showed weaker ROS signals in OE plants and stronger accumulation in KD lines, consistent with lower H O levels and higher SOD activity in OE compared to KD plants (Figure 7d, e). Collectively, these results demonstrate that *MaBBX21* enhances heat tolerance by promoting flavonol accumulation, suppressing anthocyanin biosynthesis, and maintaining cellular redox balance through efficient regulation of ROS levels.

### 2.9 *MaBBX21* Enhances UV-B Tolerance by Promoting Flavonoid Biosynthesis and ROS Scavenging

BBX proteins play important roles in signaling cascades under diverse abiotic stresses. To examine the role of MaBBX21 under UV-B stress, two-month-old WT, *MaBBX21*-OE, and KD plants were exposed to UV-B irradiation. After 72 h of treatment, no visible lesions were observed in either OE or KD plants compared to WT, indicating the absence of severe morphological damage under the tested conditions (Figure S20a). At the transcriptional level, UV-B exposure induced the expression of *MaBBX21, MaWRKY23, MaGCN5, MaFLS1, MaDFR2,* and *MaANS* in WT plants, whereas their expression was reduced in KD lines (Figure S20b).

Metabolite profiling revealed enhanced accumulation of flavonoid compounds, including quercetin, kaempferol, rutin, as well as anthocyanin derivatives (cyanidin and delphinidin) in OE lines, while their levels were reduced in KD plants relative to WT (Figure S20c). This suggests that MaBBX21 positively regulates UV-B–induced flavonoid biosynthesis. Furthermore, improved stress tolerance in OE lines was associated with better redox homeostasis. Histochemical staining (DAB and NBT) showed weaker ROS accumulation in OE plants and stronger signals in KD lines, consistent with lower H O content and higher SOD activity in OE compared to KD plants (Figure S21a-b). Overall, these findings indicate that MaBBX21 enhances UV-B tolerance by promoting flavonoid (both flavonol and anthocyanin) accumulation and maintaining cellular redox balance through efficient regulation of ROS levels.

## 3 Discussion

Flavonoids in land plants function not only as determinants of pigmentation and nutritional value but also as essential protective metabolites that mitigate abiotic stress through their antioxidant and photoprotective properties (Davies et al. 2022). Banana is a globally important crop for both nutrition and economic value, and heat-induced yield instability threatens food security and livelihoods, particularly in developing countries where it serves as a staple and key export commodity (Varma et al 2019). In the widely cultivated Grand Naine (AAA) banana cultivar, anthocyanin accumulation is predominantly restricted to the light exposed bract, whereas flowers, enclosed within the bract, remain largely colorless, reflecting a tightly controlled spatial regulation of flavonoid biosynthesis. To decipher the differential pigmentation patterns, we performed comparative transcriptome analysis of anthocyanin-rich bracts, colorless flowers and most environmentally exposed leaves. This analysis revealed a strong enrichment of light-responsive transcription factors and flavonoid biosynthesis genes in bracts and leaves. In banana, we identified MaBBX21 as a previously uncharacterized central regulatory node that integrates transcriptional activation via *MaWRKY23* and epigenetic modulation via *MaGCN5* to coordinate tissue-specific and stress-responsive flavonoid accumulation in banana.

BBX transcription factors are well-established regulators of plant development and environmental signalling, particularly through their roles in light perception, photomorphogenesis, and specialized metabolism (Xu et al. 2016; Fu et al. 2024; Gómez-Ocampo et al. 2024). Their involvement in anthocyanin biosynthesis is well known across plant species. For instance, MdBBX21 enhances light-induced anthocyanin accumulation in apple fruit peel, while BBX16 activates MYB10 to promote pigmentation in red pear (Bai et al. 2019b). Comparable regulatory functions have been reported in tomato, sweet cherry, citrus, poplar, and apple, where BBX proteins fine-tunes anthocyanin biosynthesis in response to environmental cues such as light, temperature and UV radiation (Fang et al. 2019; Li et al. 2021; Luo et al. 2023; Fu et al. 2024). Consistent with these observations, our results demonstrate that MaBBX21 functions as a positive regulator of anthocyanin accumulation in banana which was visually observed in the OE lines. The MaBBX21 interacts with MaHY5, a central light-responsive transcription factor, forming a conserved MaBBX–MaHY5 regulatory module. This module and MaBBX21 independently facilitates activation of key anthocyanin biosynthetic genes, including *MaDFR2* and *MaANS,* thereby promoting pigment accumulation.

Beyond its role in anthocyanin biosynthesis, MaBBX21 also exerts significant influence over flavonol accumulation as evident by high flavonol accumulation in OE lines and reduced in KD lines, indicating its broader role in modulating flavonoid metabolic flux. Flavonol synthase (FLS1) catalyzes the formation of major flavonols such as quercetin, kaempferol, and myricetin (Cao et al. 2024). The flavonols biosynthesis is primarily controlled by SG7 R2R3-MYB transcription factors, including AtMYB11, AtMYB12, and AtMYB111 in *Arabidopsis thaliana* (Mehrtens et al. 2005; Pandey et al. 2015; Li et al. 2019). However, our previous findings (Naik et al. 2025) indicate that this canonical SG7 MYB module plays a moderate role in banana. In this study, we found that MaBBX21 does not directly activate the *MaFLS1* promoter by dual-luciferase assays, and the expression of SG7 MYBs was slightly changed in *MaBBX21*-OE and KD lines. Therefore, suggesting the involvement of other intermediate regulators in flavonol biosynthesis.

WRKY transcription factors are increasingly recognized as versatile regulators of specialized metabolism, functioning as either activators or repressors depending on the regulatory context (Zhang et al. 2023). For example, AtWRKY23 in *Arabidopsis thaliana* and NtWRKY11 in *Nicotiana tabacum* positively regulate flavonol biosynthesis, whereas VvWRKY70 represses flavonol accumulation in *Vitis vinifera* (Grunewald et al. 2012; Wang et al. 2021; Wei et al. 2023). We selected homologs of flavonol biosynthesis-related WRKY TFs in banana and found significantly higher expression of *MaWRKY23* in *MaBBX21*-OE lines and downregulation in KD lines. Further, our findings demonstrate that MaBBX21 directly binds to and activates the *MaWRKY23* promoter. In turn, MaWRKY23 activates *MaFLS1,* thereby promoting flavonol biosynthesis while simultaneously exerting a moderate repressive effect on the anthocyanin branch. Collectively, these results uncover a previously unrecognized BBX–WRKY regulatory module that fine-tunes the balance between anthocyanin and flavonol biosynthesis.

In addition to transcriptional regulation, our study reveals a critical epigenetic layer underlying flavonoid biosynthesis. Transcription factors can modulate chromatin architecture by recruiting or activating histone-modifying enzymes (Candela-Ferre et al. 2024). Several TF–chromatin regulator interactions have been reported, including AtWRKY40–AtJMJ17, AtWRKY53–AtHDA9, AtBES1–AtREF6/ELF6, AtPIF7–AtREF6, and transcriptional activation of AtREF6 by AtHSFA2 (Yu et al. 2008; Liu et al. 2019; Zheng et al. 2020; Wang et al. 2021; Cheng et al. 2024). Histone acetylation, particularly H3K9ac, is strongly associated with active transcription in light-responsive pathways, including those regulated by HY5 and HYH (Charron et al. 2009). A few studies linking H3K9ac to activation of flavonoid biosynthesis genes are known in Arabidopsis and Petunia (Li et al. 2020; Patrick et al. 2021). GCN5 is a conserved HAT that promotes gene expression through deposition of H3K9/14 acetylation marks and facilitates RNA polymerase II recruitment (Vlachonasios et al. 2003; Hu et al. 2015). However, its involvement in secondary metabolism has remained largely unresolved. Here, we demonstrate that MaBBX21 directly activates *MaGCN5* transcription. This regulatory relationship is supported by transactivation assays and *in vivo* binding analyses and is accompanied by a substantial increase in global H3K9ac levels in *MaBBX21*-OE plants. Importantly, MaBBX21 promotes H3K9 acetylation at promoters of key flavonoid biosynthesis genes, including *MaANS* and *MaDFR,* via MaGCN5. Furthermore, MaGCN5 acetylates the *MaBBX21* promoter itself, suggesting the presence of a positive feedback loop that reinforces pathway activation and stabilizes gene expression. These findings provide direct mechanistic evidence linking transcription factor activity with chromatin remodeling and redefine flavonoid biosynthesis as an epigenetically reinforced regulatory program rather than a purely transcription-driven process.

The regulatory role of MaBBX21 extends beyond metabolism to encompass abiotic stress adaptation. BBX transcription factors are increasingly recognized as integrators of environmental signals and redox homeostasis (Wang et al. 2013; Xu et al. 2020). For instance, MdBBX37 enhances cold tolerance in apple by activating ROS detoxification pathways, while AtBBX18, AtBBX19, and OsBBX4 regulate tolerance to heat, drought, and salinity through ABA- and heat shock factor–mediated transcriptional pathways (Wang et al. 2013; Xu et al. 2020; An et al. 2021). Similarly, SlBBX17 promotes thermotolerance in tomato, and IbBBX24 enhances drought and salinity tolerance in sweet potato through ROS-scavenging mechanisms (Xu et al. 2022; Zhang et al. 2022). Consistent with these observations, *MaBBX21*-OE banana plants exhibit enhanced tolerance to heat and UV-B stress in contrast to WT and KD lines. Further, MaBBX21-KD lines were more sensitive to these stresses as compare to WT. This enhanced stress tolerance appears to result from coordinated metabolic and epigenetic reprogramming. GCN5 plays a key role in abiotic stress responses by promoting histone acetylation and activating stress-responsive genes such as *HSFA3* and *UVH6* (Hu et al. 2015). In agreement with this, *MaGCN5* is induced under heat stress, suggesting its epigenetic regulation in heat stress mitigation.

During heat stress, the expression of *MaFLS1* was markedly induced, resulting in enhanced accumulation of flavonols such as kaempferol, myricetin, and rutin in MaBBX21-OE lines, whereas KD lines exhibited the opposite trend. In contrast, the expression of *MaDFR2* and *MaANS* declined at later stages of stress, coinciding with reduced anthocyanin accumulation. This shift in metabolic flux suggests a preferential channeling toward flavonol biosynthesis under heat stress conditions. Flavonols are well known for their strong antioxidant capacity and role in mitigating heat-induced oxidative damage, thereby contributing to stress tolerance (Naik et al. 2024; Agati et al. 2012). Additionally, the presence of pre-accumulated anthocyanins in MaBBX21-OE lines may provide basal antioxidant protection, reducing the need for sustained anthocyanin biosynthesis during prolonged stress. In the case of UV-B stress, the expression of both flavonol- and anthocyanin-biosynthetic genes was induced, along with increased accumulation of kaempferol, rutin, delphinidin, and cyanidin in *MaBBX21*-OE lines, in contrast to KD lines. This coordinated increase suggests a protective metabolic response, where flavonols function as UV-B screening compounds and anthocyanins act as antioxidants to scavenge ROS, thereby enhancing stress tolerance (Agati et al. 2012). Metabolic flux may be redirected toward other protective pathways, including polyamine biosynthesis, lipid and wax metabolism, heat shock protein activation, and carbohydrate metabolism (Li et al. 2022; Maryam et al. 2025). Heat and UV-B stresses impose distinct physiological demands, leading to differential regulation of flavonoid metabolism. Under heat stress, metabolic flux is preferentially redirected toward flavonol biosynthesis, as flavonols provide efficient and rapidly deployable antioxidant protection for maintaining redox homeostasis, while the anthocyanin branch is often transcriptionally repressed and energetically costly, resulting in reduced accumulation. In contrast, UV-B stress requires both UV screening and antioxidative defense, promoting the accumulation of both flavonols and anthocyanins.

In conclusion, this study establishes a multilayered regulatory framework in which MaBBX21 functions as a central integrator of light signaling, transcriptional regulation, and epigenetic modulation to control flavonoid biosynthesis in banana. Through coordinated interactions with MaHY5, MaWRKY23, and the histone acetyltransferase MaGCN5, MaBBX21 promotes locus-specific enrichment of activating chromatin marks such as H3K9ac, thereby establishing a permissive transcriptional environment at key biosynthetic genes. This regulatory module enables precise control of metabolic flux between flavonol and anthocyanin branches and highlights MaBBX21 as a dynamic regulator of pathway balance. The integration of light signaling with chromatin remodeling represents a novel mechanism linking environmental cues to stable metabolic outputs in secondary metabolism. Furthermore, the tight association between enhanced flavonoid accumulation and improved stress tolerance underscores the dual nutritional and adaptive significance of this pathway (Figure 8). Future studies focusing on anthocyanin accumulation in fruit pulp of OE and KD lines will further evaluate the translational potential of this regulatory module. Overall, this work provides a robust conceptual and mechanistic framework for engineering nutritionally enriched and climate-resilient banana cultivars.

**FIGURE 8.**
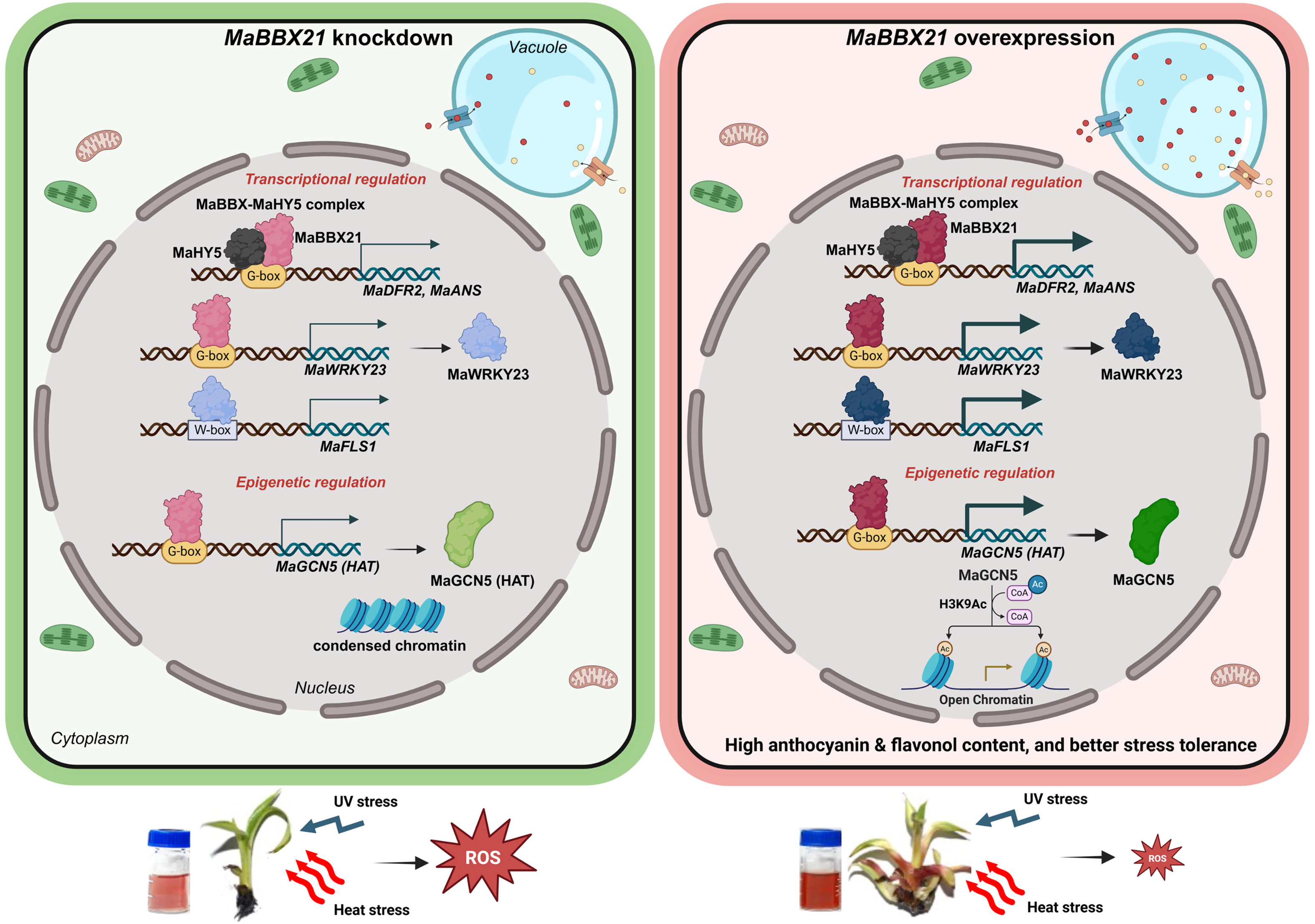
Proposed working model illustrating how MaBBX21 integrates transcriptional and epigenetic regulation to control flavonoid biosynthesis in banana. MaBBX21 functions both independently and in association with MaHY5, forming a MaBBX21–MaHY5 complex that activates the expression of *MaDFR2* and *MaANS* by binding to BBX-binding motifs in their promoter regions. Consistent with this regulatory role, the expression of these key anthocyanin biosynthesis genes was significantly reduced in *MaBBX21*-KD plants. In contrast, *MaBBX21*-OE lines showed markedly increased expression of *MaDFR2* and *MaANS,* resulting in visibly enhanced anthocyanin accumulation in *MaBBX21*-OE plants. Furthermore, MaBBX21 was found to transcriptionally activate *MaWRKY23*, which subsequently binds to the W-box element in the promoter of *MaFLS1*, thereby promoting flavonol accumulation. This regulatory module fine-tunes metabolic flux within the flavonoid pathway by enhancing *MaFLS1* expression while limiting flux toward *MaDFR2*, thus maintaining a balanced allocation between flavonol and anthocyanin biosynthesis. In parallel, MaBBX21 directly activates the histone acetyltransferase *MaGCN5*, which increases H3K9 acetylation at the promoter regions of *MaDFR2* and *MaANS* near the transcription start site (TSS), thereby reinforcing anthocyanin production. Additionally, MaGCN5 enhances H3K9 acetylation at the promoter of *MaBBX21* itself near the TSS, suggesting the presence of a positive feedback regulatory mechanism. Collectively, these integrated transcriptional and epigenetic regulatory mechanisms promote flavonoid accumulation. Notably, the elevated flavonoid content enhances reactive oxygen species (ROS) scavenging capacity, thereby conferring improved tolerance to heat and UV-B stress in banana.

## 4 Material and Methods

### 4.1 Plant Materials and Growth Conditions

*Musa acuminata* cv. Grand Naine (AAA genome) plants were cultivated in the experimental field of the BRIC-National Institute of Plant Genome Research (NIPGR), New Delhi. Planting materials were obtained from the NIPGR field facility. Embryogenic cell suspension (ECS) cultures were established from immature male flower buds of the Grand Naine cultivar and used for stable transgenic generation and ChIP assays. Six- to eight-week-old Grand Naine saplings were used for heat and UV-B treatments. Plants were maintained under controlled growth chamber conditions (Model: AR-41L3, Percival Scientific USA) at 26 °C with a 16 h light/8 h dark photoperiod, light intensity of 250 μmol m ² s ¹, and 60% relative humidity. *Nicotiana benthamiana* seeds were germinated on Murashige and Skoog (MS) medium at 24 °C under a 16 h light/8 h dark photoperiod with a light intensity of 100 μmol m ² s ¹ for one week. Seedlings were subsequently transplanted into a soil:vermiculite mixture (1:1, v/v) and grown in a greenhouse at 22 °C under a 16 h photoperiod for five weeks prior to experimentation.

### 4.2 RNA Isolation and RNA-seq Analysis of Bract, Flower, And Leaf Tissues

Bract, flower, and leaf tissues were harvested and immediately frozen in liquid nitrogen to preserve RNA integrity. Total RNA was extracted using the Spectrum™ Plant Total RNA Kit (Sigma-Aldrich, Darmstadt, Germany) following the manufacturer’s protocol. RNA purity and concentration were measured using a NanoDrop™ 2000 spectrophotometer (Thermo Fisher Scientific, Waltham, MA, USA), while integrity was assessed by agarose gel electrophoresis and confirmed using an Agilent 2100 Bioanalyzer (Agilent Technologies, Santa Clara, CA, USA). Residual genomic DNA was removed using the TURBO DNA-free™ Kit (Invitrogen, Waltham, MA, USA). Only high-quality RNA samples (RIN ≥ 7) from three biological replicates per tissue were used for RNA-seq library construction. Paired-end libraries were prepared using the KAPA mRNA HyperPrep Kit (Illumina-compatible) and assessed on an Agilent 4150 TapeStation system. Sequencing was performed on the Illumina NovaSeq 6000 platform, and library quality was further validated using HSD100 ScreenTape®. After removing adapters and low-quality reads, clean reads were aligned to the *Musa acuminata* reference genome from the Banana Genome Hub using STAR (v2.7.10a) (Dobin et al. 2013). Transcript assembly and quantification were conducted with StringTie (v2.2.1) (Pertea et al. 2015), and resulting read counts were used for differential expression analysis with edgeR (Robinson et al. 2010). Log -transformed TPM values were used for heatmap visualization. Gene Ontology and pathway enrichment analyses were performed using AgriGO v2.0 and g:Profiler, respectively (Reimand et al. 2016; Tian et al. 2017).

### 4.3 RT-qPCR Gene Expression Analysis

First-strand cDNA was synthesized from 1 µg of DNase-treated RNA using the RevertAid H Minus First Strand cDNA Synthesis Kit (Thermo Fisher Scientific) with oligo(dT) primers. RT-qPCR was performed on a 7500 Fast Real-Time PCR System (Applied Biosystems, Waltham, MA, USA) using SYBR Green chemistry. Each 10 µL reaction included 2× SYBR Green PCR Master Mix, gene-specific primers, and cDNA equivalent to 10 ng total RNA. The *MaACTIN1* gene (GenBank: AF246288) was used as an internal reference, and relative expression levels were calculated using the 2^–ΔΔCT method (Livak & Schmittgen, 2001). Results are presented as fold changes relative to the control or lowest-expressing tissue.

### 4.4 Identification of Flavonoids Regulators and Gene Cloning

Phylogenetic analyses were conducted for *MaBBX* protein to investigate their evolutionary relationships. Protein sequences were aligned using the MUSCLE algorithm implemented in MEGA 11 software (Tamura et al. 2021). Phylogenetic trees were constructed using the Maximum Likelihood (ML) method with default substitution models. Statistical support for each node was evaluated by bootstrap analysis with 1000 replicates. The resulting trees were visualized and annotated using Interactive Tree of Life (iTOL v7N) (Letunic & Bork, 2021).

Full-length CDSs of *MaBBX20*, *MaBBX21*, *MaHY5*, *MaWRKY23*, and *MaGCN5* (without stop codons) were amplified from banana leaf cDNA using gene-specific primers containing Gateway™ attB sites, designed from Banana Genome Hub sequences. High-fidelity PCR products were cloned into pDONR™Zeo (Invitrogen) via BP recombination and transformed into *Escherichia coli* DH5α, with positive clones confirmed by Sanger sequencing.

### 4.5 Transient OE and Gene KD in Banana Fruit Discs

For transient OE, the CDSs of *MaBBX21*, *MaWRKY23*, and *MaGCN5* were cloned into the Gateway-compatible vector pANIC6b, which enables strong monocot expression in monocots driven by ZmUBI promoter. The vector contains a proOsACT1-driven hygromycin resistance marker, a proPvUBI-driven GUSPlus reporter. Constructs and empty vector (EV) controls were transformed into *A. tumefaciens* EHA105 (Mehra et al. 2019; Rajput et al. 2025) and introduced into banana fruit slices by vacuum infiltration. Following 3 days of co-cultivation on modified MS medium, transformation efficiency was confirmed by GUS staining. For gene KD, 250–300 bp fragments of *MaBBX21* and *MaGCN5* were cloned into pANIC8b (Mann et al. 2012), transformed into EHA105, and infiltrated into fruit discs and ECS under similar conditions. Transformed tissues were harvested in liquid nitrogen, stored at –80 °C, and used for RT-qPCR and targeted flavonoid/metabolite profiling by LC–MS.

### 4.6 Generation of *MaBBX21* Overexpression and Knockdown Transgenic Lines

Stable transgenic lines of banana were generated via Agrobacterium-mediated genetic transformation. ECS were initiated from the immature male flowers of the Grand Nain cultivar following the protocol described by (Tripathi et al. 2015). Banana ECSs was transformed with the *A. tumefaciens* strain EHA105 harbouring either the OE construct or the KD construct of the *MaBBX21* gene. The transformation procedure was carried out according to the protocol developed by (Tiwari, 2019). Following co-cultivation, the transformed ECS were selected on selection medium supplemented with 25 mg/L hygromycin. Hygromycin-resistant putative transformants were further screened and confirmed further by a GUS histochemical assay. The transgene expression level was subsequently analysed by RT-qPCR. The confirmed transgenic lines were transferred to multiplication medium containing 2.5 mg/L 6-benzylaminopurine (BAP) and 0.2 mg/L indole-3-acetic acid (IAA) to promote shoot proliferation. Well-developed shoots were then transferred to an MS rooting medium supplemented with 2 mg/L indole-3-butyric acid (IBA) to induce root formation.

### 4.7 Histochemical **β**-Glucuronidase (GUS) Assay

Transgenic plant tissues were incubated in an X-gal containing (GUS) staining solution at 37 °C overnight according to Saxena et al, 2023. After staining, the samples were cleared of chlorophyll and excess stain by washing several times with, and then documented.

### 4.8 DPBA Staining and Total Anthocyanin Quantification

DPBA staining was performed to visualize flavonol accumulation (quercetin and kaempferol) in roots of 15-day-old banana seedlings (WT, *MaBBX21*-OE, and KD lines). Root segments (∼6 mm) were incubated in 0.5% DPBA (in methanol) for 15 min in the dark, briefly rinsed, and observed under a fluorescence/confocal microscope using UV excitation (365–405 nm). Total anthocyanin content was quantified following Singh et al. (2024) with minor modifications. Briefly, 25 mg dry tissue was extracted in acidified methanol (1% v/v) overnight at 28 °C in the dark, centrifuged, and absorbance was measured at 530 and 657 nm. Anthocyanin content was calculated as (A −0.25 × A) per gram fresh weight.

### 4.9 LC-MS For Detection of Targeted Flavonoids

Targeted flavonoid profiling was performed following established protocols (Pandey et al. 2016; Saxena et al. 2023) with minor modifications. To differentiate glycosylated and aglycone forms, both non-hydrolyzed and acid-hydrolyzed extracts were prepared. For non-hydrolyzed samples, powdered tissue was extracted in 80% methanol, incubated at 70 °C for 15 min, centrifuged (12,000 × g, 10 min), and the supernatant was collected. For hydrolyzed samples, the extract was treated with 1 N HCl (1:3, v/v), incubated at 94 °C for 2 h, followed by ethyl acetate extraction. The organic phase was dried under reduced pressure (90–100 mbar, 37 °C) using a Rotavapor R-300 (BUCHI, Flawil, Switzerland), reconstituted in 80% methanol, and filtered (0.22 µm) prior to LC–MS analysis. Anthocyanin quantification was carried out as described by Singh et al. (2024). Lyophilized samples (200 mg) were extracted in 80% methanol, filtered (0.22 µm), and subjected to acid hydrolysis using 2 M methanol:HCl (3:1, v/v) at 90 °C for 45 min. Samples were dried, resuspended in 80% methanol, and analyzed using an Exion LC UPLC system (Sciex) coupled to a QTRAP 6500+ mass spectrometer (AB Sciex) with electrospray ionization.

### 4.10 Promoter Isolation and Reporter Construct Preparation

Promoter sequences of target genes were retrieved from the Banana Genome Hub based on *Musa acuminata* genome annotations (D’hont et al. 2012). Cis-regulatory elements were analyzed using the New PLACE database with manual validation of key motifs. Genomic DNA was isolated from banana leaf tissue using the DNeasy Plant Mini Kit (Qiagen, Hilden, Germany).

Promoter regions (*proMaGCN5*-1.7 kb, *proMaWRKY23*-1.7 kb, *proMaFLS1*-2.5 kb, *proMaDFR2*-1.5 kb, and *proMaANS*-1.2 kb) were amplified using gene-specific primers and Phusion™ High-Fidelity DNA Polymerase (Thermo Fisher Scientific), and cloned into the pENTR™/D-TOPO™ vector. Positive clones were confirmed by Sanger sequencing. Verified entry clones were recombined into the p635nRRF vector containing the pro35S::REN cassette for normalization (Kumar et al. 2018) via Gateway® LR reaction to generate dual-luciferase reporter constructs.

### 4.11 Yeast Two-Hybrid and Yeast One[Hybrid Assay

For protein–protein interaction analysis, CDSs of *MaBBX20*, *MaBBX21*, and *MaHY5* were cloned into the yeast two-hybrid vectors pGADT7g (prey) and pGBKT7g (bait) (Clontech). Constructs were co-transformed into *Saccharomyces cerevisiae* Y2HGold using the lithium acetate method and selected on SD/-Trp/-Leu medium at 30 °C. Positive colonies were assayed on stringent SD/-Trp/-Leu/-His/-Ade medium supplemented with X-Gal (20 µg/ml) and 3-AT (50 mM). Interactions were determined based on yeast growth and blue color development. The p53–SV40 large T antigen interaction served as a positive control (Pipas & Levine, 2001), while empty vector combinations were used as negative controls.

For yeast one-hybrid (Y1H) assays, promoter fragments of *MaFLS1*, *MaDFR2*, *MaANS*, *MaWRKY23*, and *MaGCN5* were cloned into the pABAi vector, integrated into Y1HGold yeast (Takara Bio Inc.) following the Matchmaker™ Gold protocol. CDSs of *MaBBX21* and *MaWRKY23* were cloned into pGADT7-GW and transformed into promoter-integrated strains with EV controls. Minimum inhibitory concentrations (MIC) of Aureobasidin A (AbA) were optimized for each promoter: 1 µM (*MaFLS1*), 900 nM (*MaDFR2*), 850 nM (*MaANS*), 900 nM (*MaFLS1* W-box), 600 nM (*MaWRKY23*), and 400 nM (*MaGCN5* F1/F2). Protein–DNA interactions were confirmed by growth on AbA-containing selective medium.

### 4.12 Subcellular Localization and Bimolecular Fluorescence Complementation (BiFC) Assays

Entry clones of *MaBBX20, MaBBX21,* and *MaGCN5* were recombined into the pSITE-3CA vector to generate N-terminal YFP fusion constructs, following the method described by Chakrabarty et al. (2007), with the empty vector used as a control. The resulting constructs were introduced into *Agrobacterium tumefaciens* GV3101-pMP90 and resuspended in infiltration buffer (10 mM MgCl, 10 mM MES-KOH, pH 5.7, and 200 µM acetosyringone), followed by incubation for 3 h. Equal volumes of nYFP and cYFP fusion cultures were mixed in a 1:1 ratio and co-infiltrated into *Nicotiana benthamiana* leaves, with empty vector combinations serving as controls.

For *in planta* protein–protein interaction analysis using BiFC, the coding sequences (CDSs) of MaBBX21 and MaHY5 were recombined into the 35S-pSITE-nYFP-N1 and 35S-pSITE-cYFP-N1 vectors, respectively, as described by Martin et al. (2009). After incubation for 48 h at 22 °C in the dark, YFP fluorescence was detected with excitation at 514 nm and emission at 527 nm, while RFP fluorescence was detected with excitation at 561 nm and emission collected between 580–620 nm using a Leica TCS SP5 confocal laser-scanning microscope (Leica Microsystems, Wetzlar, Germany).

### 4.13 Split and Dual Luciferase Assay

For *in planta* protein–protein interaction analysis, the CDS of *MaHY5* was cloned into pCAMBIA1300-nLUC and *MaBBX21* into pCAMBIA1300-cLUC to generate split luciferase fusion constructs (Chen et al. 2008). Constructs were transformed into *Agrobacterium tumefaciens* GV3101, resuspended in infiltration buffer and co-infiltrated into *N. benthamiana* leaves. After 48 h, leaves were treated with D-luciferin (100 µM), and luciferase activity was detected using a CCD imaging system (Bio-Rad).

For transactivation assays, CDSs of *MaBBX21*, *MaHY5*, and *MaWRKY23* were cloned into pSITE-3CA under the 2×35S promoter (Chakrabarty et al. 2007). Promoters of *MaFLS1*, *MaDFR2*, *MaANS*, *MaWRKY23*, and *MaGCN5* were fused to LUC in p635nRRF, with REN as internal control. Effector and reporter constructs were co-infiltrated into *N. benthamiana* (Walter et al. 2004), and after 48 h, LUC/REN activity was measured using the Dual-Luciferase® Reporter Assay System (Promega). Results are presented as mean ± SD of four biological replicates.

### 4.14 ChIP-qPCR Assay in Banana Embryonic Cell Suspension

Chromatin immunoprecipitation (ChIP) assays were performed following Saleh et al. (2008) with modifications for banana ECS. *MaBBX21-GFP*, *MaWRKY23-GFP*, and *MaGCN5-GFP* constructs were transiently expressed via 30 min vacuum infiltration. At 4 days post-infiltration, ∼3 g tissue was harvested, cross-linked with 1% formaldehyde, frozen, and ground. Nuclei were isolated, and chromatin was sheared using a Bioruptor Plus (Diagenode) to obtain 200–500 bp fragments. Chromatin was pre-cleared with Protein A agarose beads and immunoprecipitated using anti-GFP (Abcam, ab290) or anti-H3K9ac (Abcam, ab10812), with anti-IgG (Abcam, ab48386) as a negative control. After elution and reverse cross-linking, DNA was purified. ChIP-qPCR was performed using SYBR Green Master Mix (Applied Biosystems), and enrichment was calculated using the 2^−ΔCT method (Livak & Schmittgen, 2001) relative to IgG controls.

### 4.15 Protein Isolation and Immunoblotting

For *in vivo* acetylase activity assays, total proteins were extracted from *MaBBX21-*OE and WT banana leaves were ground in liquid nitrogen and homogenized in cold protein extraction buffer as described in Ahmed et al. 2025. The clarified supernatants were subjected to immunoblot analysis using appropriate antibodies. Histone acetylation levels were assessed using an anti-H3K9ac antibody. Ponceau S staining of Rubisco was used as a loading control for protein normalization.

### 4.16 HAT Inhibitor Treatment

One-month-old *MaBBX21*-OE, KD, and WT banana plants were treated with α-methylene-γ-butyrolactone (Cat no. 226416 SIGMA) under controlled growth conditions. Leaf samples were harvested at the 12 h time point, and the expression levels of target genes were analyzed by RT-qPCR.

### 4.17 Heat and UV-B Treatment to Banana Saplings

Two-month-old *in vitro*-grown WT and *MaBBX21*-OE banana plants of comparable size were used for stress treatment. Leaf samples were collected at 0 h, 24 h, 48 h and 72 h after the onset of stress, immediately frozen in liquid nitrogen, and stored at -80 °C until further analysis. Stress experiments were conducted in a controlled plant growth chamber (Model: AR-41L3; Percival, Perry, IA, USA) under a 16 h light /8 h dark photoperiod, light intensity of 250 μmol m ² s ¹, and 60% relative humidity. Heat stress was imposed by exposing plants to 42 °C, while control plants were maintained at 26 °C. UV-B irradiation was applied using two UV-B light sources (TL 20W/01 RS SLV/25) positioned at a distance of 40 cm from the plants. For phenotypic representation, photographs were captured at the 72 h time point under continuous heat and UV-B stress, showing both WT plants and three independent *MaBBX21*-OE lines.

### 4.18 NBT and DAB Staining, H[O[ Quantification, and SOD Activity Assays

For heat and UV-B stress analysis, the third fully expanded leaf from the apex was harvested. Reactive oxygen species (ROS) were detected histochemical using NBT staining for superoxide (·O) and DAB staining for H O (Daudi & O’Brien, 2012). For NBT staining, leaves were vacuum-infiltrated with 0.1 mg ml ¹ NBT in 25 mM HEPES buffer (pH 7.8), incubated in the dark at 25 °C for 2 h, and decolorized in 80% ethanol. H O content was quantified following Alexieva et al. (2001) by reacting leaf extracts (in 0.1% TCA) with potassium phosphate buffer and KI, and measuring absorbance at 390 nm against a standard curve. For enzymatic assays, leaf tissue (200 mg) was homogenized in extraction buffer (50 mM PB pH 7.5, 1 mM PMSF, 2% PVPP, 0.1 mM EDTA) and centrifuged (16,000 × g, 20 min, 4 °C). The supernatant was used for superoxide dismutase (SOD) activity, with protein quantified by the Bradford method (Bradford, 1976). SOD activity was measured based on inhibition of NBT photoreduction in a reaction containing riboflavin, methionine, EDTA, and NBT, with absorbance recorded at 560 nm. One unit of SOD activity was defined as the amount of enzyme causing 50% inhibition of NBT reduction (Tyagi et al. 2024).

### 4.19 Statistical Analysis

Statistical analyses were conducted using one-way and two-way analysis of variance (ANOVA), followed by Dunnett’s and Sidak’s multiple comparisons test, respectively. For specific pairwise comparisons, unpaired Student’s t-tests were applied. Statistical significance is indicated as follows: P ≤ 0.05 (*), P ≤ 0.01 (**), P ≤ 0.001 (***), P ≤ 0.0001 (****); ns denotes non-significant differences. The statistical analysis was performed using GraphPad Prism 8.0 software (GraphPad Software, Boston, MA, USA).

## Authors’ Contributions

SS and AP conceived the idea and designed the research. ST & BP performed transcriptome analysis. RN & SB performed genetic transformation of banana. SS, ST, HC and JN conducted the experiments. SS, ST, and AP interpreted the data. SS & ST wrote the manuscript. SS, ST, RS, and AP edited and finalized the manuscript. AP coordinated the research project. All authors read and approved the final manuscript.

## Acknowledgements

This work was supported by the core grant of BRIC-National Institute of Plant Genome Research and Department of Science and Technology (DST)- Anusandhan National Research Foundation (ANRF) for Core Research Grant (CRG/2022/001178) to AP. SS & HC acknowledge University Grant Commission Fellowship, Government of India, for Senior Research Fellowships. ST & SB acknowledge DST, ANRF for National Post Doctoral Fellowship (N-PDF). The authors are thankful to DBT-eLibrary Consortium (DeLCON) for providing access to e-resources. We acknowledge the Metabolome and Confocal facility at BRIC-NIPGR for phytochemical and imaging analysis, respectively.

## Data Availability Statement

All data supporting the findings of this study are available within the paper and within the supplemental data published online. The raw sequencing data generated in this study have been deposited in the NCBI SRA under the Bio-Project accession number PRJNA1449240.

## Supporting Information

**Figure S1:** Flavonoids content in bract, flower and leaf tissues. (a) Bar graphs depicting the differential accumulation of key aglycon flavonols (quercetin and myricetin), glycan flavonols (rutin and kaempferol 3-O-rutinoside), dihydromyricetin and naringenin in bract, flower and leaf tissue. (b) Differential accumulation of different anthocyanidin derivatives in bract, flower and leaf tissue. (c) Quantification of proanthocyanidins (catechin, procyanidin B2, procyanidin C1 and epigallocatechin) in bract flower and leaf tissue. Asterisks represent statistically significant differences (\**p* < 0.05, \*\**p* < 0.01, \*\*\**p* < 0.001, \*\*\*\**p* < 0.0001, ns = not significant).

**Figure S2:** Expression profiling of flavonoid biosynthesis genes. (a) Pictorial representation of flavonoids biosynthesis pathway along with the heatmaps showing the log2 normalized TPM expression value of biosynthesis genes involved at different steps. Colour bar scale (green to magenta) indicate low expression to high expression. (b) Relative expression analysis of different flavonoids biosynthesis pathway genes by RT-qPCR in bract, flower and leaf tissues. Standard deviation and significance level are represented by error bars and asterisks, respectively. Asterisks represent statistically significant differences (\**p* < 0.05, \*\**p* < 0.01, \*\*\**p* < 0.001, \*\*\*\**p* < 0.0001, ns = not significant).

**Figure S3:** DEG distribution and venn diagram analysis among bract, flower, and leaf tissues. (a) The bar plot shows the total number of significantly upregulated (blue) and downregulated (pink) differentially expressed genes (DEGs) identified in three pairwise comparisons: bract vs flower (B vs F), bract vs leaf (B vs L), and flower vs leaf (F vs L). (b) Venn diagrams showing the unique and overlapping sets of differentially expressed genes (DEGs) among pairwise comparisons of upregulated DEGs and downregulated DEGs identified in B vs F, B vs L, and F vs L comparisons.

**Figure S4:** Gene Ontology (GO) enrichment analysis of differentially expressed genes (DEGs) in bract, flower, and leaf tissues. Bubble plots show GO enrichment results for upregulated (↑) and downregulated (↓) DEGs identified in bract (B), flower (F), and leaf (L) tissues in biological process: (a) B (↑) vs F (↓); (b) F (↑) vs B (↓); (c) F (↑) vs L (↓); (d) L (↑) vs F (↓); (e) B (↑) vs L (↓); (f) L (↑) vs B (↓). The x-axis represents fold enrichment of each GO term. Bubble size corresponds to the number of DEGs associated with each term, and colour intensity reflects the statistical significance by -log10 *p*-value.

**Figure S5:** KEGG pathway enrichment analysis of DEGs in bract, flower, and leaf tissues. Bubble plots depict KEGG pathway enrichment of upregulated (↑) and downregulated (↓) DEGs across bract (B), flower (F), and leaf (L) tissues: (a) B (↑) vs F (↓); (b) B (↓) vs F (↑); (c) B (↑) vs L (↓); (d) B (↓) vs L (↑); (e) F (↑) vs L (↓); (f) F (↓) vs L (↑). The x-axis indicates fold enrichment of each pathway, representing the degree of overrepresentation. Bubble size reflects the number of DEGs involved in each pathway, and the color-scale denotes adjusted *p*-value significance.

**Figure S6:** Expression profiling of light signalling and WRKY transcription factor family genes. (a) Heatmap showing the differential expression patterns of key light signalling-related transcription factor genes including *MaBBX,* and (b) *MaCOP*, *MaPIFs*, *MaSPA* and *MaCRY* in bract, flower, and leaf tissues, revealing distinct tissue-specific transcriptional regulation. c Heatmap showing the expression *MaWRKY* transcription factor genes, and in bract, flower and leaf tissue. The color scale represents log -normalized gene expression values (green and magenta reflects low and high expression levels, respectively).

**Figure S7:** Phylogenetic analysis and subcellular localization of MaBBXs in banana. (a) Phylogenetic tree of BBX proteins from banana and selected plant species, illustrating their clustering into five distinct groups (I to V). Group II, III and IV were furthered divided into different subgroups. MaBBX20 (Ma01_g03030) and MaBBX21 (Ma07_g18870) of banana clusters within subgroup iv of group IV and shows very close homology to previously characterized AtBBX20, AtBBX21 and OsBBX21, which are known regulators of anthocyanin biosynthesis. Branch lengths are indicated along each branch. Bootstrap support values are visualized as periwinkle-colored bubbles, with bubble size proportional to the level of support at each node. (b) Multiple sequence alignment of BBX proteins from subgroup IV-iv, showing conserved residues within two B-box zinc finger-like motifs, indicated by black stars. The presence of two conserved B-box zinc finger-like motifs is a characteristic feature of subgroup IV, where they are involved in protein–protein interactions. (c) Expression trends of MaBBX genes belonging to subgroup iv of group IV were further validated by RT-qPCR analysis in bract, flower and leaf tissue. Statistical significance is indicated by asterisks: *P* ≤ 0.05 (\**), P* ≤ *0.01 (*\*\**), P* ≤ *0.001 (*\*\*\*), *P* ≤ 0.0001 (****); ns = not significant. (d) Subcellular localization of MaBBX20 and MaBBX21 in *N. benthamiana* leaves. YFP-tagged MaBBX20 and MaBBX21 fusion proteins localize to the nucleus, co-localizing with the nuclear marker NLS-RFP, as observed via confocal microscopy. The EV (pSITE-3CA) expressing free YFP was used as a negative control. Scale bar = 50 µm.

**Figure S8:** Expression analysis in transient *MaBBX20*-OE in banana fruit discs. Expression analysis of *MaBBX20, MaDFR1/2,* and *MaANS* in banana fruit slices of *MaBBX20-*OE and EV-OE. Statistical significance is indicated by asterisks: *P* ≤ 0.05 (\**), P* ≤ *0.01 (*\*\**), P* ≤ *0.001 (*\*\*\*), *P* ≤ 0.0001 (****); ns = not significant.

**Figure S9:** Expression analysis and metabolite profiling in transient *MaBBX21*-OE and KD banana fruit discs. (a) Representative GUS-stained images of banana fruit discs demonstrate successful transient transformation with *MaBBX21*-OE and KD constructs. The total anthocyanin quantification showed visible difference in anthocyanin content in EV, *MaBBX21*-OE and *MaBBX21*-KD fruit discs. Images were captured 3 d post-transformation. (b) Relative expression levels of *MaBBX21* gene and flavonoid biosynthesis genes *(MaCHS3–6, MaCHI1/2, MaF3H1/2, MaF3’H, MaF3’5’H1/7*, *MaFLS1/2, MaDFR1/2,* and *MaANS)* were quantified by RT-qPCR in *MaBBX21-*OE and *MaBBX21*-KD banana fruit slices. Expression profiles showed differential modulation compared to corresponding EV controls. (c) Quantification of flavonol (myricetin, kaempferol) and anthocyanin derivatives (cyanidin, petunidin, delphinidin, peonidin) in *MaBBX21* OE and *MaBBX21*-KD fruit discs. Statistical significance is indicated by asterisks: *P* ≤ 0.05 (\**), P* ≤ *0.01 (*\*\**), P* ≤ *0.001 (*\*\*\*), *P* ≤ 0.0001 (****); ns = not significant.

**Figure S10:** Screening and validation of stable *MaBBX21*-OE and KD transgenic lines. (a) Expression analysis of *MaBBX21* in overexpression lines determined by RT–qPCR. (b) Expression analysis of *MaBBX21* in KD lines determined by RT–qPCR. (c) Representative GUS-stained images of banana leaves confirming successful stable transformation with the *MaBBX21*-OE lines and d *MaBBX21*-KD lines. Statistical significance is indicated by asterisks: *P* ≤ 0.05 (\**), P* ≤ *0.01 (*\*\**), P* ≤ *0.001 (*\*\*\*), *P* ≤ 0.0001 (****); ns = not significant.

**Figure S11:** Expression analysis of flavonoid biosynthesis genes in *MaBBX21*-OE and KD lines Expression analysis of *MaFLS2, MaF3’H,* and *MaDFR1* in (a) *MaBBX21*-OE lines and (b) *MaBBX21*-KD lines compared with WT plants. Statistical significance is indicated by asterisks: *P* ≤ 0.05 (\**), P* ≤ *0.01 (*\*\**), P* ≤ *0.001 (*\*\*\*), *P* ≤ 0.0001 (****); ns = not significant.

**Figure S12:** Flavonoids content in *MaBBX21*-OE and KD lines. (a) The levels of flavonols (kaempferol, quercetin, myricetin), rutin and anthocyanins including cyanidin, delphinidin, peonidin, petunidin and pelargonidin were altered in *MaBBX21*-OE and (b) KD lines relative to the control WT plants. Statistical significance is indicated by asterisks: *P* ≤ 0.05 (\**), P* ≤ *0.01 (*\*\**), P* ≤ *0.001 (*\*\*\*), *P* ≤ 0.0001 (****); ns = not significant.

**Figure S13:** Protein–protein interaction analysis of MaBBX20 and MaHY5. Interaction analysis of MaBBX20 and MaHY5 using (a) yeast two-hybrid assay and (b) bimolecular fluorescence complementation showing no detectable physical interaction between the two proteins.

**Figure S14:** Expression analysis of *MaBBX21* downstream regulatory gene. (a) Expression analysis of flavonoid regulatory genes, including *MaMYB*s (*MaMYBAN1/2, MaMYBPA1/2, MaMYBFA1/2/3), MabHLH*s *(MabHLH2/7/9), MaWD40s* (*MaWD40.1/2*), and *MaWRKY*s (*MaWRKY32/42/50*) in *MaBBX21-*OE lines, and (b) *MaBBX21*-KD lines by RT–qPCR. Statistical significance is indicated by asterisks: *P* ≤ 0.05 (\**), P* ≤ *0.01 (*\*\**), P* ≤ *0.001 (*\*\*\*), *P* ≤ 0.0001 (****); ns = not significant.

**Figure S15:** Transient overexpression of *MaWRKY23* in banana fruit slices. (a) Expression analysis of early flavonoid biosynthesis genes *MaCHS1–6, MaCHI1–2, MaF3H1–2, MaFLS2, MaF3’H, MaF3*′*5*′*H1–7* and *MaDFR1* by RT–qPCR showing modulated expression in *MaWRKY23*-OE banana fruit slices. (b) Quantification of anthocyanidins (peonidin and pelargonidin) in transient overexpression of *MaWRKY23* showing their accumulation levels in comparison to EV control. Statistical significance is indicated by asterisks: *P* ≤ 0.05 (\**), P* ≤ *0.01 (*\*\**), P* ≤ *0.001 (*\*\*\*), *P* ≤ 0.0001 (****); ns = not significant.

**Figure S16:** Expression analysis of MaBBX21 downstream *HAT* genes. (a) Relative expression of histone acetyltransferase genes *HAG2* and *HAG3* in *MaBBX21*-OE and *MaBBX21*-KD banana lines compare to the control (WT) by RT–qPCR. Statistical significance is indicated by asterisks: *P* ≤ 0.05 (\**), P* ≤ *0.01 (*\*\**), P* ≤ *0.001 (*\*\*\*), *P* ≤ 0.0001 (****); ns = not significant. (b) Subcellular localization of MaGCN5 in *N. benthamiana* leaves showed that the MaGCN5–YFP fusion protein localized to the nucleus and co-localized with the nuclear marker NLS-RFP, as observed by confocal microscopy. Scale bar = 30 µm.

**Figure S17:** Expression analysis in *MaGCN5*-OE and KD banana fruit discs. (a) Bar graphs showing the relative expression of *MaWRKY23* and flavonol biosynthesis genes (*MaFLS1/2, MaF3’H*, and *MaDFR1)* in *MaGCN5-OE* and KD in banana fruit discs and (b) ECS, as determined by RT–qPCR relative to their respective EV controls. Statistical significance is indicated by asterisks: *P* ≤ 0.05 (\**), P* ≤ *0.01 (*\*\**), P* ≤ *0.001 (*\*\*\*), *P* ≤ 0.0001 (****); ns = not significant.

**Figure S18:** Occupancy of MaGCN5 on the promoters of *MaDFR2, MaANS,* and *MaBBX21* determined using anti-GFP antibody. Enrichment of MaGCN5 on the promoters of *MaDFR2, MaANS,* and *MaBBX21* at specific fragments, as detected by ChIP–qPCR using an anti-GFP antibody. Statistical significance is indicated by asterisks: *P* ≤ 0.05 (\**), P* ≤ *0.01 (*\*\**), P* ≤ *0.001 (*\*\*\*), *P* ≤ 0.0001 (****); ns = not significant.

**Figure S19:** *MaBBX21* enhances heat stress tolerance by modulating flavonoid biosynthesis. (a) Relative expression analysis of *MaWRKY23*, *MaGCN5* and *MaANS* and (b) LCMS based quantification of flavonoid (rutin, quercetin and delphinidin) in the control (WT), *MaBBX21*-OE and KD lines at 0 h, 24 h, 48 h and 72 h of heat stress treatment. Statistical significance is indicated by asterisks: *P* ≤ 0.05 (\**), P* ≤ *0.01 (*\*\**), P* ≤ *0.001 (*\*\*\*), *P* ≤ 0.0001 (****); ns = not significant.

**Figure S20:** MaBBX21 confers UV-B tolerance by enhancing flavonoid biosynthesis and limiting ROS accumulation. (a) Phenotypic comparison of 45 d old saplings of WT and *MaBBX21*-OE and KD lines grown under normal conditions and subsequently exposed to UV-B stress for 72 h. (b) Expression analysis of *MaBBX21, MaWRKY23, MaGCN5, MaFLS1, MaDFR2* and *MaANS*; and (c) quantification of different flavonoids in WT, *MaBBX21*-OE and KD lines at 0 h, 24 h, 48 h, and 72 h of UV-B treatment. Data represent mean ± SD of WT and three independent OE lines, each with three technical replicates. Statistical significance is indicated by asterisks: *P* ≤ 0.05 (\**), P* ≤ *0.01 (*\*\**), P* ≤ *0.001 (*\*\*\*), *P* ≤ 0.0001 (****); ns = not significant.

**Figure S21.** Staining of ROS and quantification of H O content and SOD enzyme activity. (a) DAB and NBT staining in the leaves of WT and *MaBBX21*-OE and KD lines after 48 h of UV-B treatment. (b) Estimation of H O content and SOD enzyme activity in WT, *MaBBX21*-OE and KD lines at 48 h of UV-B stress. Standard deviation and significance level are represented by error bars and asterisks, respectively. Asterisks represent statistically significant differences (\**p* < 0.05, \*\**p* < 0.01, \*\*\**p* < 0.001, \*\*\*\**p* < 0.0001, ns = not significant).

