## Supplementary Figures for "A MaBBX21-MaGCN5 Histone Acetyltransferase Module Regulates Flavonoid Biosynthesis to Improve Heat and UV-B Stress Tolerance in Banana (*Musa acuminata*)"

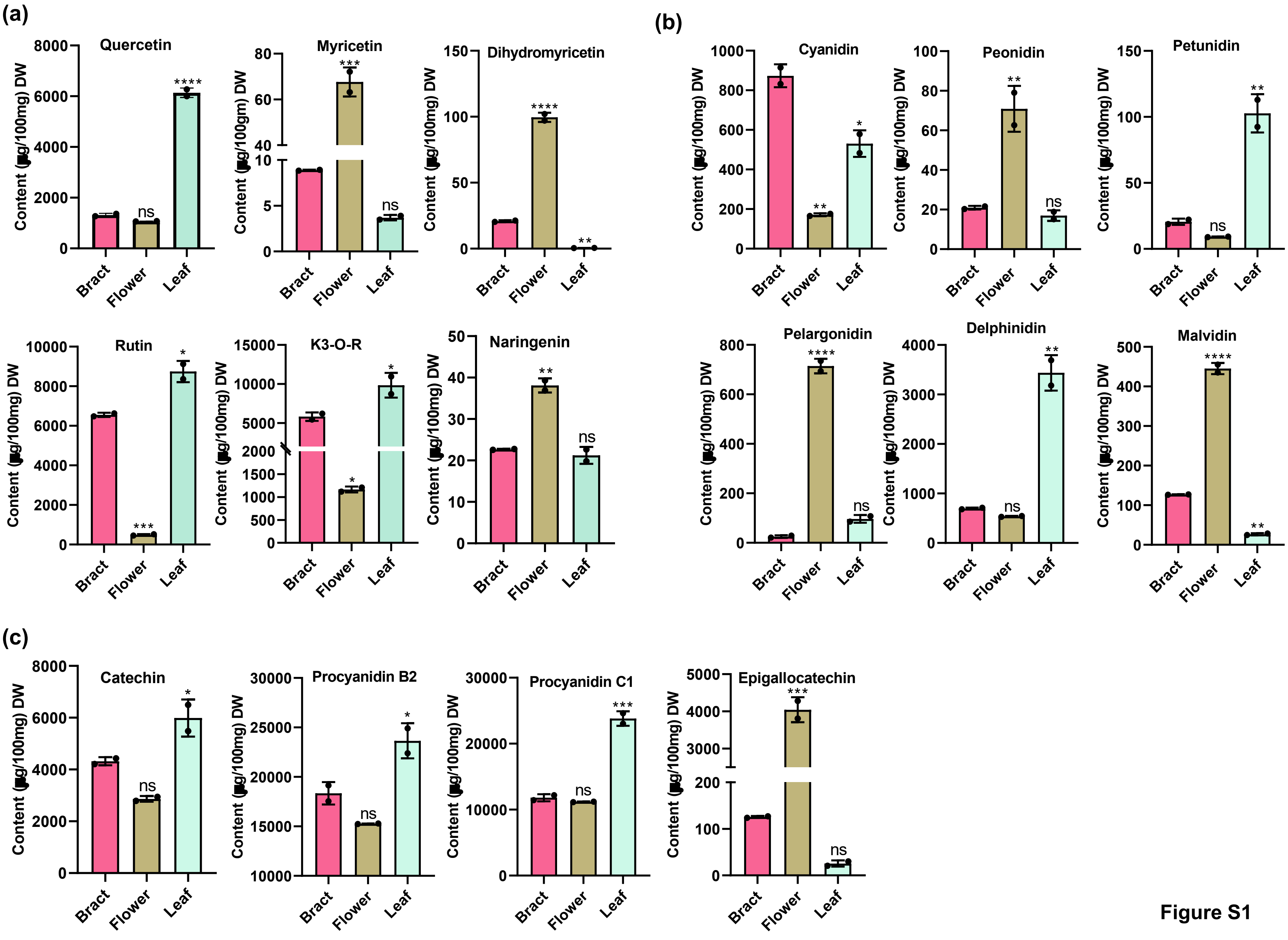
(a)
Quercetin
Myricetin
Dihydromyricetin
Rutin
K3-O-R
Naringenin
(b)
Cyanidin
Peonidin
Petunidin
Pelargonidin
Delphinidin
Malvidin
(c)
Catechin
Procyanidin B2
Procyanidin C1
Epigallocatechin
Bract
Flower
Leaf
ns

Figure S1

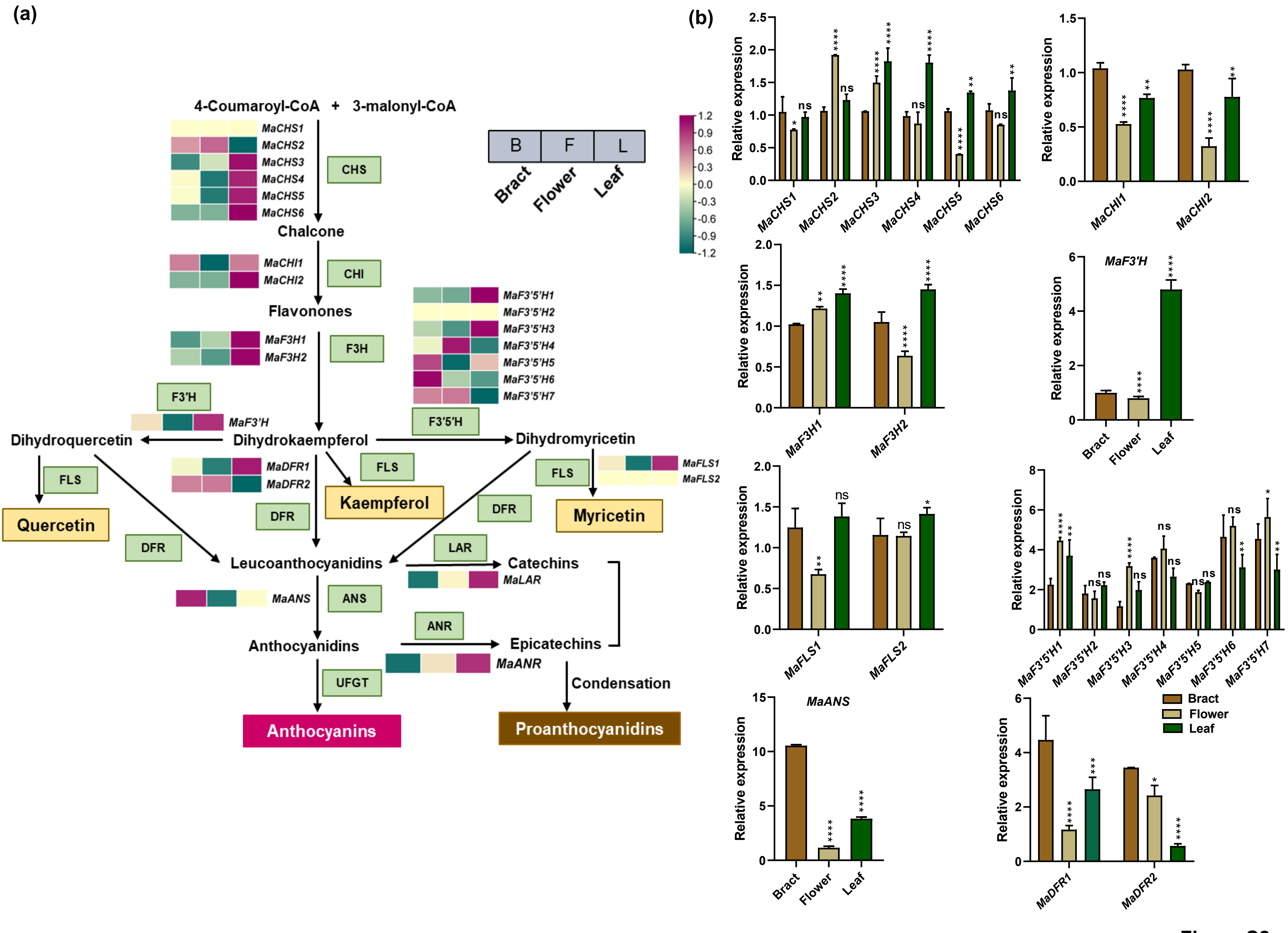

(a)
4-Coumaroyl-CoA + 3-malonyl-CoA
MaCHS1
MaCHS2
MaCHS3
MaCHS4
MaCHS5
MaCHS6
CHS
Chalcone
MaCHI1
MaCHI2
CHI
Flavonones
MaF3H1
MaF3H2
F3H
F3'H
MaF3'H
Dihydroquercetin
Dihydrokaempferol
Dihydromyricetin
MaF3'5'H1
MaF3'5'H2
MaF3'5'H3
MaF3'5'H4
MaF3'5'H5
MaF3'5'H6
MaF3'5'H7
F3'5'H
B F L
Bract Flower Leaf
FLS
Quercetin
MaDFR1
MaDFR2
Kaempferol
MaFLS1
MaFLS2
Myricetin
DFR
LAR
Leucoanthocyanidins
Catechins
MaLAR
MaANS
ANS
ANR
Anthocyanidins
Epicatechins
MaANR
UFGT
Condensation
Anthocyanins
Proanthocyanidins
(b)
Relative expression
MaCHS1 MaCHS2 MaCHS3 MaCHS4 MaCHS5 MaCHS6
MaCHI1 MaCHI2
MaF3H1 MaF3H2
MaF3'H
MaFLS1 MaFLS2
MaF3'5'H1 MaF3'5'H2 MaF3'5'H3 MaF3'5'H4 MaF3'5'H5 MaF3'5'H6 MaF3'5'H7
MaANS
MaDFR1 MaDFR2
Bract
Flower
Leaf


Figure S2

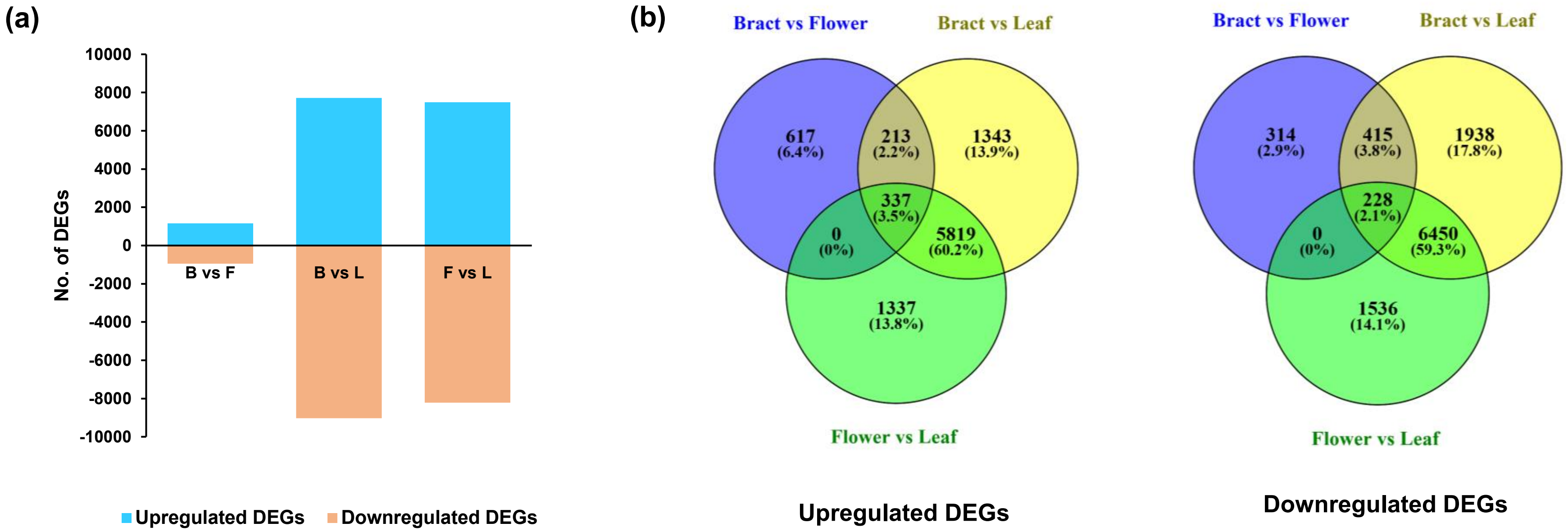

(a)
No. of DEGs
10000
8000
6000
4000
2000
0
-2000
-4000
-6000
-8000
-10000
B vs F
B vs L
F vs L
Upregulated DEGs
Downregulated DEGs
(b)
Bract vs Flower
Bract vs Leaf
Flower vs Leaf
617 (6.4%)
213 (2.2%)
1343 (13.9%)
337 (3.5%)
0 (0%)
5819 (60.2%)
1337 (13.8%)
Upregulated DEGs
Bract vs Flower
Bract vs Leaf
Flower vs Leaf
314 (2.9%)
415 (3.8%)
1938 (17.8%)
228 (2.1%)
0 (0%)
6450 (59.3%)
1536 (14.1%)
Downregulated DEGs


Figure S3

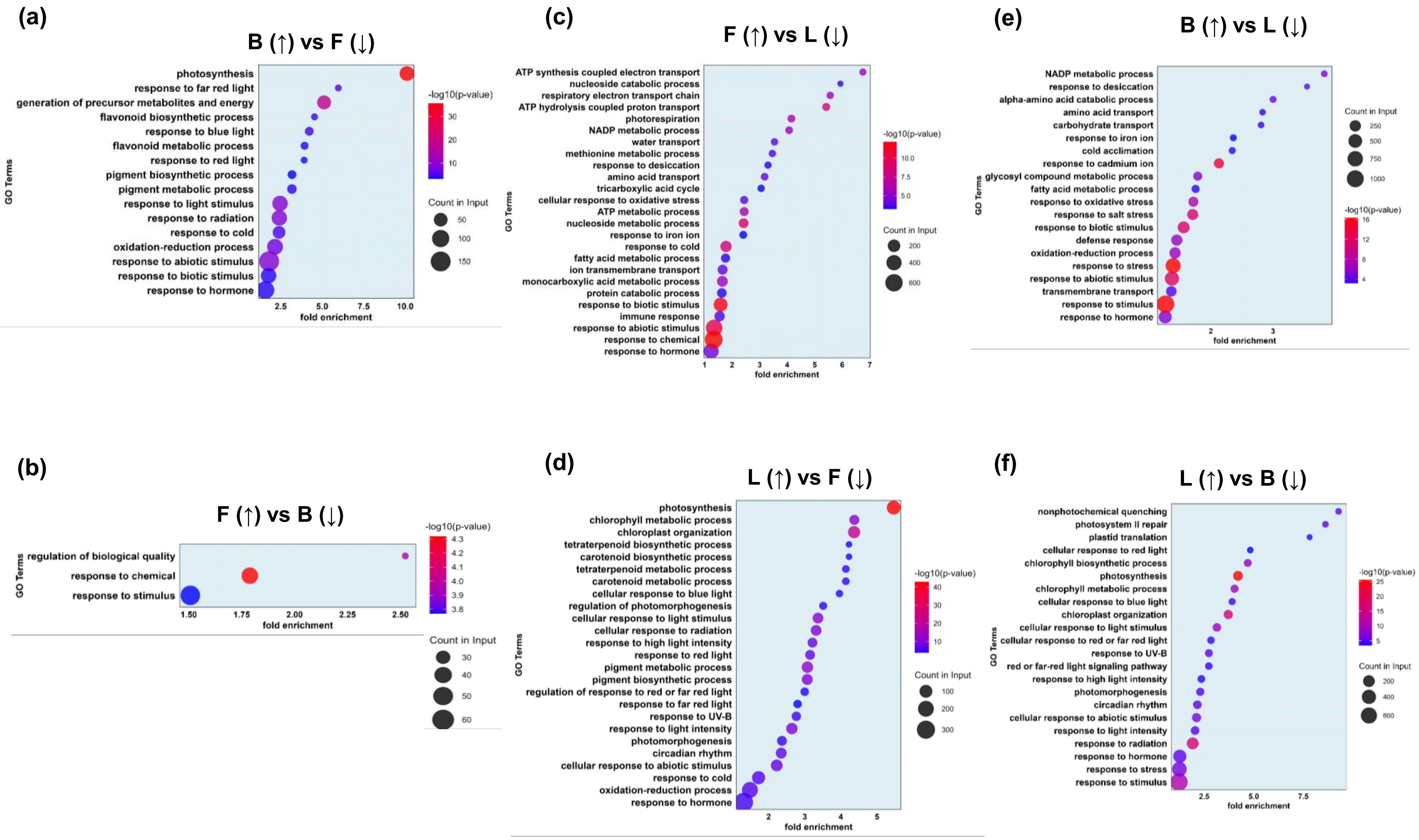

(a)
B (↑) vs F (↓)
photosynthesis
response to far red light
generation of precursor metabolites and energy
flavonoid biosynthetic process
response to blue light
flavonoid metabolic process
response to red light
pigment biosynthetic process
pigment metabolic process
response to light stimulus
response to radiation
response to cold
oxidation-reduction process
response to abiotic stimulus
response to biotic stimulus
response to hormone
GO Terms
fold enrichment
-log10(p-value)
Count in Input
(b)
F (↑) vs B (↓)
regulation of biological quality
response to chemical
response to stimulus
GO Terms
fold enrichment
-log10(p-value)
Count in Input
(c)
F (↑) vs L (↓)
ATP synthesis coupled electron transport
nucleoside catabolic process
respiratory electron transport chain
ATP hydrolysis coupled proton transport
photorespiration
NADP metabolic process
water transport
methionine metabolic process
response to desiccation
amino acid transport
tricarboxylic acid cycle
cellular response to oxidative stress
ATP metabolic process
nucleoside metabolic process
response to iron ion
response to cold
fatty acid metabolic process
ion transmembrane transport
monocarboxylic acid metabolic process
protein catabolic process
response to biotic stimulus
immune response
response to abiotic stimulus
response to chemical
response to hormone
GO Terms
fold enrichment
-log10(p-value)
Count in Input
(d)
L (↑) vs F (↓)
photosynthesis
chlorophyll metabolic process
chloroplast organization
tetraterpenoid biosynthetic process
carotenoid biosynthetic process
tetraterpenoid metabolic process
carotenoid metabolic process
cellular response to blue light
regulation of photomorphogenesis
cellular response to light stimulus
cellular response to radiation
response to high light intensity
response to red light
pigment metabolic process
pigment biosynthetic process
regulation of response to red or far red light
response to far red light
response to UV-B
response to light intensity
photomorphogenesis
circadian rhythm
cellular response to abiotic stimulus
response to cold
oxidation-reduction process
response to hormone
GO Terms
fold enrichment
-log10(p-value)
Count in Input
(e)
B (↑) vs L (↓)
NADP metabolic process
response to desiccation
alpha-amino acid catabolic process
amino acid transport
carbohydrate transport
response to iron ion
cold acclimation
response to cadmium ion
glycosyl compound metabolic process
fatty acid metabolic process
response to oxidative stress
response to salt stress
response to biotic stimulus
defense response
oxidation-reduction process
response to stress
response to abiotic stimulus
transmembrane transport
response to stimulus
response to hormone
GO Terms
fold enrichment
Count in Input
-log10(p-value)
(f)
L (↑) vs B (↓)
nonphotochemical quenching
photosystem II repair
plastid translation
cellular response to red light
chlorophyll biosynthetic process
photosynthesis
chlorophyll metabolic process
cellular response to blue light
chloroplast organization
cellular response to light stimulus
cellular response to red or far red light
response to UV-B
red or far-red light signaling pathway
response to high light intensity
photomorphogenesis
circadian rhythm
cellular response to abiotic stimulus
response to light intensity
response to radiation
response to hormone
response to stress
response to stimulus
GO Terms
fold enrichment
-log10(p-value)
Count in Input


Figure S4

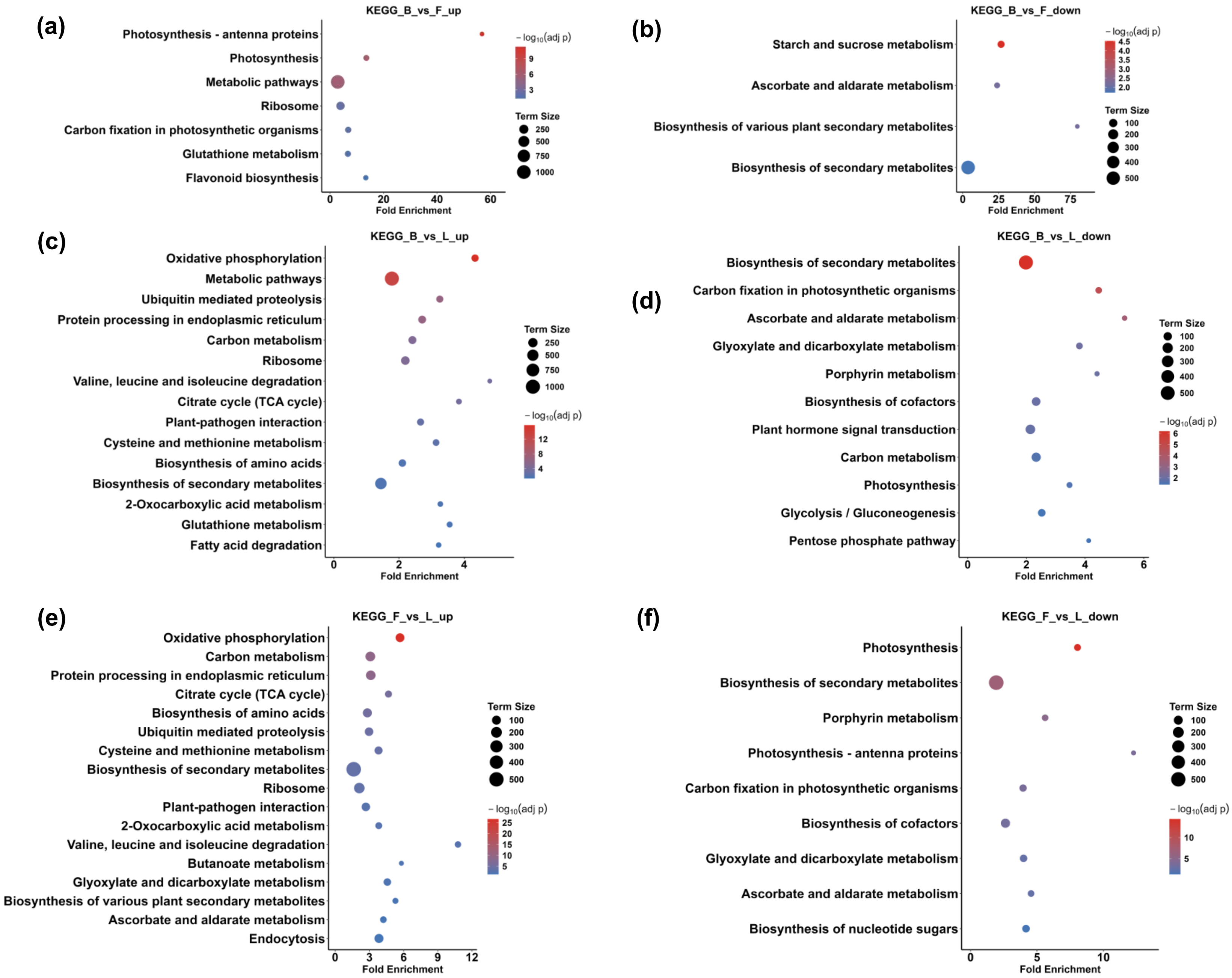

(a)
KEGG_B_vs_F_up
Photosynthesis - antenna proteins
Photosynthesis
Metabolic pathways
Ribosome
Carbon fixation in photosynthetic organisms
Glutathione metabolism
Flavonoid biosynthesis
Fold Enrichment
Term Size
(b)
KEGG_B_vs_F_down
Starch and sucrose metabolism
Ascorbate and aldarate metabolism
Biosynthesis of various plant secondary metabolites
Biosynthesis of secondary metabolites
(c)
KEGG_B_vs_L_up
Oxidative phosphorylation
Metabolic pathways
Ubiquitin mediated proteolysis
Protein processing in endoplasmic reticulum
Carbon metabolism
Ribosome
Valine, leucine and isoleucine degradation
Citrate cycle (TCA cycle)
Plant-pathogen interaction
Cysteine and methionine metabolism
Biosynthesis of amino acids
Biosynthesis of secondary metabolites
2-Oxocarboxylic acid metabolism
Glutathione metabolism
Fatty acid degradation
(d)
KEGG_B_vs_L_down
Biosynthesis of secondary metabolites
Carbon fixation in photosynthetic organisms
Ascorbate and aldarate metabolism
Glyoxylate and dicarboxylate metabolism
Porphyrin metabolism
Biosynthesis of cofactors
Plant hormone signal transduction
Carbon metabolism
Photosynthesis
Glycolysis / Gluconeogenesis
Pentose phosphate pathway
(e)
KEGG_F_vs_L_up
Oxidative phosphorylation
Carbon metabolism
Protein processing in endoplasmic reticulum
Citrate cycle (TCA cycle)
Biosynthesis of amino acids
Ubiquitin mediated proteolysis
Cysteine and methionine metabolism
Biosynthesis of secondary metabolites
Ribosome
Plant-pathogen interaction
2-Oxocarboxylic acid metabolism
Valine, leucine and isoleucine degradation
Butanoate metabolism
Glyoxylate and dicarboxylate metabolism
Biosynthesis of various plant secondary metabolites
Ascorbate and aldarate metabolism
Endocytosis
(f)
KEGG_F_vs_L_down
Photosynthesis
Biosynthesis of secondary metabolites
Porphyrin metabolism
Photosynthesis - antenna proteins
Carbon fixation in photosynthetic organisms
Biosynthesis of cofactors
Glyoxylate and dicarboxylate metabolism
Ascorbate and aldarate metabolism
Biosynthesis of nucleotide sugars


Figure S5

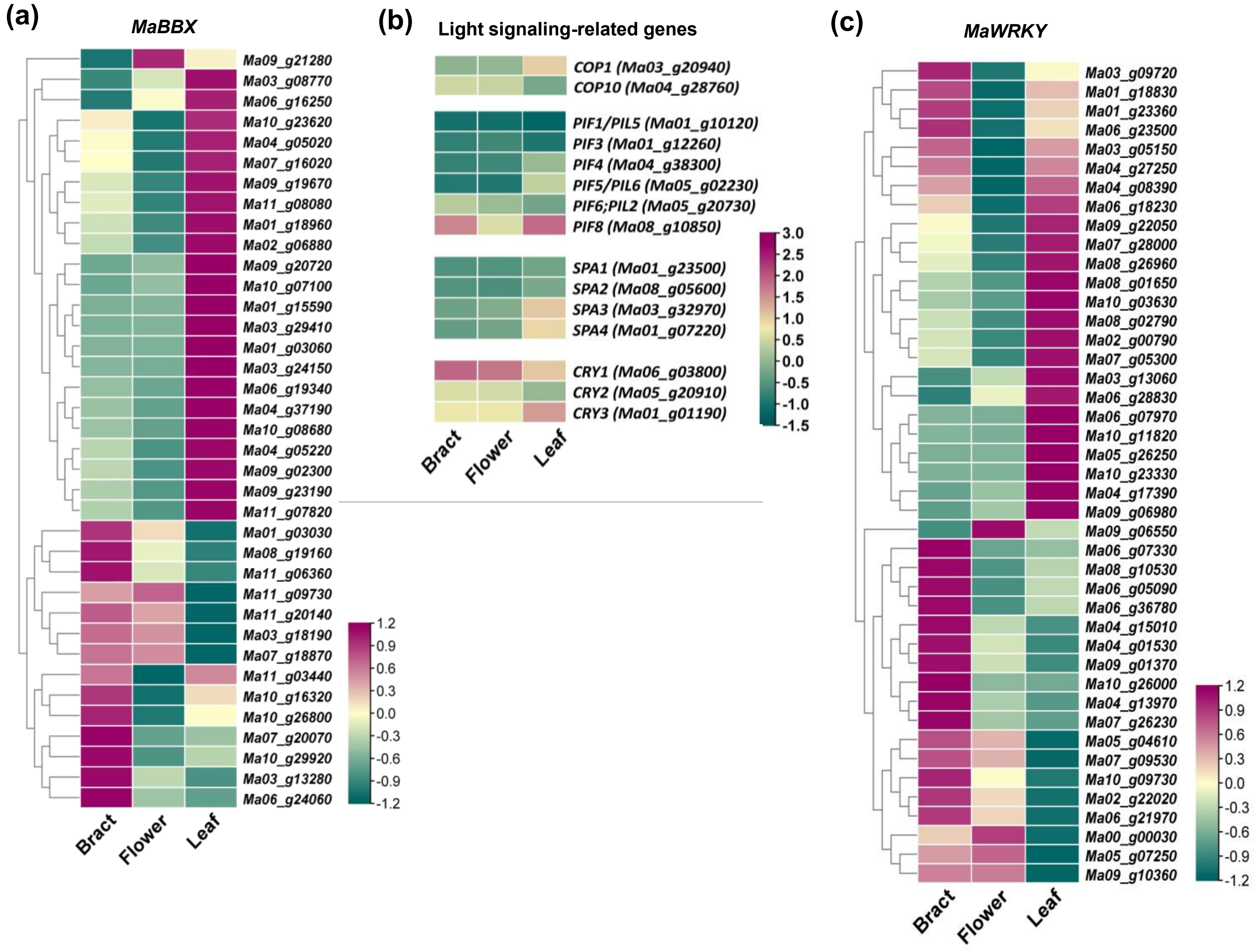
(a)
MaBBX
Ma09_g21280
Ma03_g08770
Ma06_g16250
Ma10_g23620
Ma04_g05020
Ma07_g16020
Ma09_g19670
Ma11_g08080
Ma01_g18960
Ma02_g06880
Ma09_g20720
Ma10_g07100
Ma01_g15590
Ma03_g29410
Ma01_g03060
Ma03_g24150
Ma06_g19340
Ma04_g37190
Ma10_g08680
Ma04_g05220
Ma09_g02300
Ma09_g23190
Ma11_g07820
Ma01_g03030
Ma08_g19160
Ma11_g06360
Ma11_g09730
Ma11_g20140
Ma03_g18190
Ma07_g18870
Ma11_g03440
Ma10_g16320
Ma10_g26800
Ma07_g20070
Ma10_g29920
Ma03_g13280
Ma06_g24060
1.2
0.9
0.6
0.3
0.0
-0.3
-0.6
-0.9
-1.2
Bract
Flower
Leaf
(b)
Light signaling-related genes
COP1 (Ma03_g20940)
COP10 (Ma04_g28760)
PIF1/PIL5 (Ma01_g10120)
PIF3 (Ma01_g12260)
PIF4 (Ma04_g38300)
PIF5/PIL6 (Ma05_g02230)
PIF6;PIL2 (Ma05_g20730)
PIF8 (Ma08_g10850)
SPA1 (Ma01_g23500)
SPA2 (Ma08_g05600)
SPA3 (Ma03_g32970)
SPA4 (Ma01_g07220)
CRY1 (Ma06_g03800)
CRY2 (Ma05_g20910)
CRY3 (Ma01_g01190)
3.0
2.5
2.0
1.5
1.0
0.5
0.0
-0.5
-1.0
-1.5
Bract
Flower
Leaf
(c)
MaWRKY
Ma03_g09720
Ma01_g18830
Ma01_g23360
Ma06_g23500
Ma03_g05150
Ma04_g27250
Ma04_g08390
Ma06_g18230
Ma09_g22050
Ma07_g28000
Ma08_g26960
Ma08_g01650
Ma10_g03630
Ma08_g02790
Ma02_g00790
Ma07_g05300
Ma03_g13060
Ma06_g28830
Ma06_g07970
Ma10_g11820
Ma05_g26250
Ma10_g23330
Ma04_g17390
Ma09_g06980
Ma09_g06550
Ma06_g07330
Ma08_g10530
Ma06_g05090
Ma06_g36780
Ma04_g15010
Ma04_g01530
Ma09_g01370
Ma10_g26000
Ma04_g13970
Ma07_g26230
Ma05_g04610
Ma07_g09530
Ma10_g09730
Ma02_g22020
Ma06_g21970
Ma00_g00030
Ma05_g07250
Ma09_g10360
1.2
0.9
0.6
0.3
0.0
-0.3
-0.6
-0.9
-1.2
Bract
Flower
Leaf

Figure S6

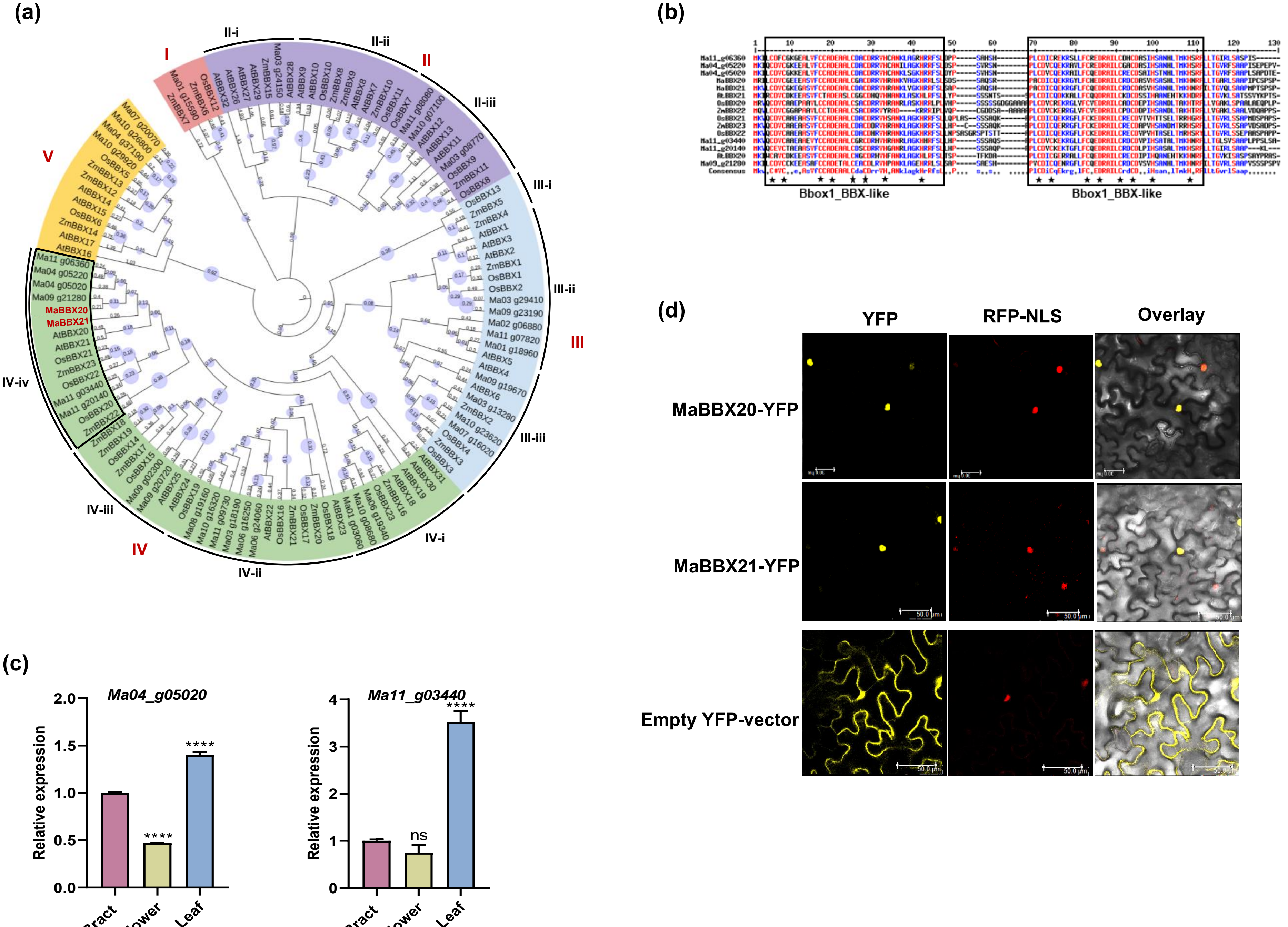

(a)
I
II
III
IV
V
II-i
II-ii
II-iii
III-i
III-ii
III-iii
IV-i
IV-ii
IV-iii
IV-iv
MaBBX20
MaBBX21
(b)
Bbox1_BBX-like
Bbox1_BBX-like
Consensus
(c)
Ma04_g05020
Ma11_g03440
Relative expression
Bract
Flower
Leaf
****
ns
(d)
YFP
RFP-NLS
Overlay
MaBBX20-YFP
MaBBX21-YFP
Empty YFP-vector
50.0 μm


Figure S7

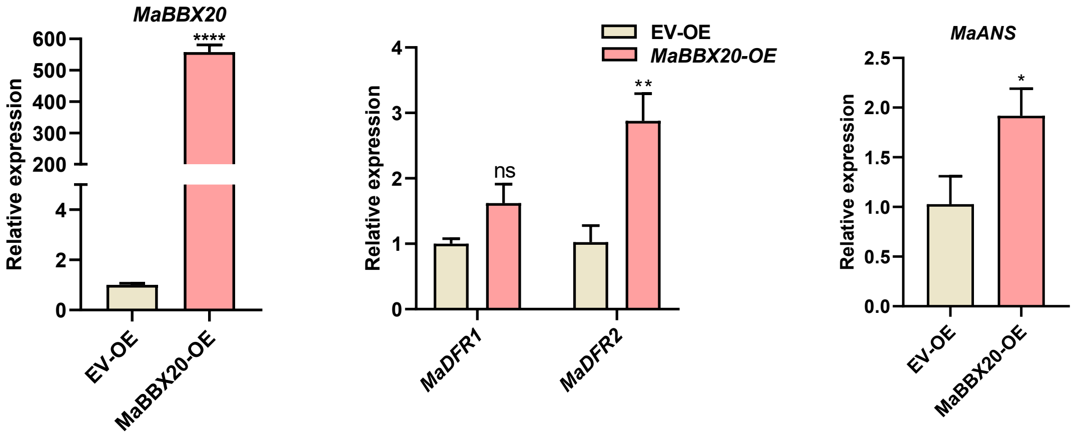

MaBBX20
Relative expression
600
500
400
300
200
4
2
0
****
EV-OE
MaBBX20-OE
EV-OE
MaBBX20-OE
4
3
2
1
0
ns
**
MaDFR1
MaDFR2
MaANS
2.5
2.0
1.5
1.0
0.5
0.0
*


Figure S8

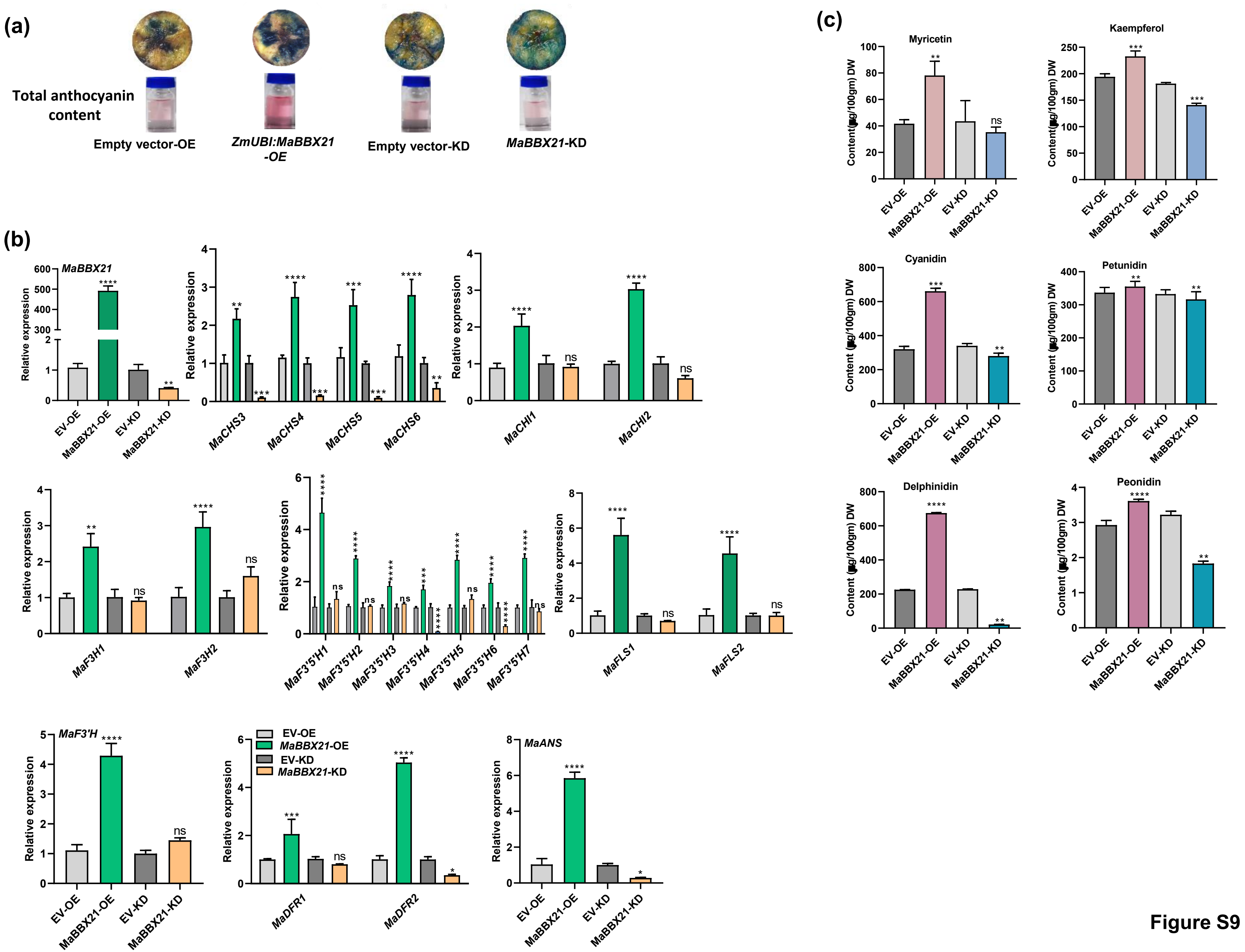

(a)
Total anthocyanin content
Empty vector-OE
ZmUBI:MaBBX21-OE
Empty vector-KD
MaBBX21-KD
(b)
MaBBX21
Relative expression
EV-OE
MaBBX21-OE
EV-KD
MaBBX21-KD
MaCHS3
MaCHS4
MaCHS5
MaCHS6
MaCHI1
MaCHI2
MaF3H1
MaF3H2
MaF3'5'H1
MaF3'5'H2
MaF3'5'H3
MaF3'5'H4
MaF3'5'H5
MaF3'5'H6
MaF3'5'H7
MaFLS1
MaFLS2
MaF3'H
MaDFR1
MaDFR2
MaANS
(c)
Myricetin
Kaempferol
Cyanidin
Petunidin
Delphinidin
Peonidin
Content(µg/100gm) DW

Figure S9

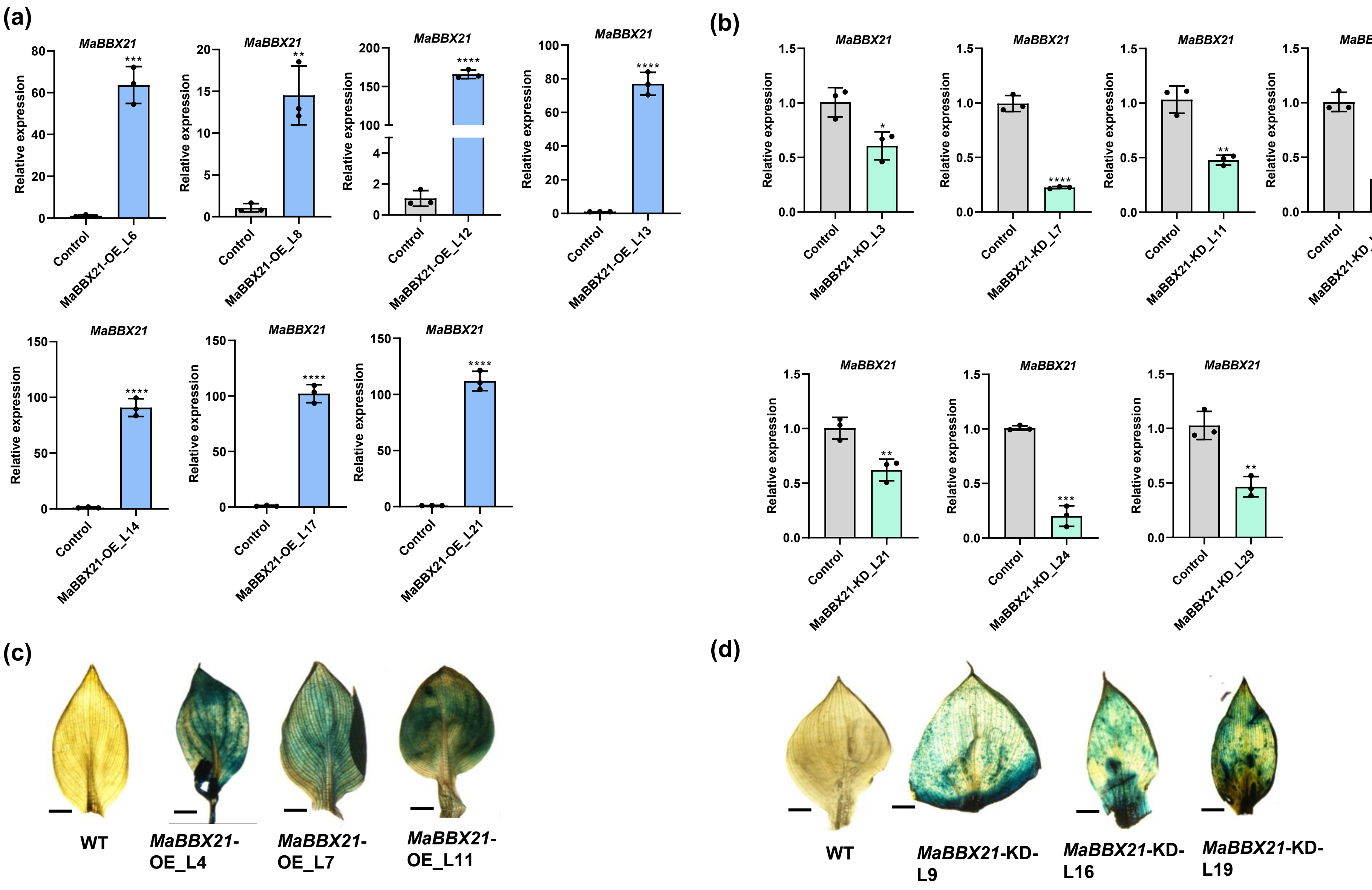

(a)
MaBBX21
Relative expression
Control
MaBBX21-OE_L6
MaBBX21-OE_L8
MaBBX21-OE_L12
MaBBX21-OE_L13
MaBBX21-OE_L14
MaBBX21-OE_L17
MaBBX21-OE_L21
(b)
MaBBX21-KD_L3
MaBBX21-KD_L7
MaBBX21-KD_L11
MaBBX21-KD_L14
MaBBX21-KD_L21
MaBBX21-KD_L24
MaBBX21-KD_L29
(c)
WT
MaBBX21-OE_L4
MaBBX21-OE_L7
MaBBX21-OE_L11
(d)
WT
MaBBX21-KD-L9
MaBBX21-KD-L16
MaBBX21-KD-L19


**Figure S10**

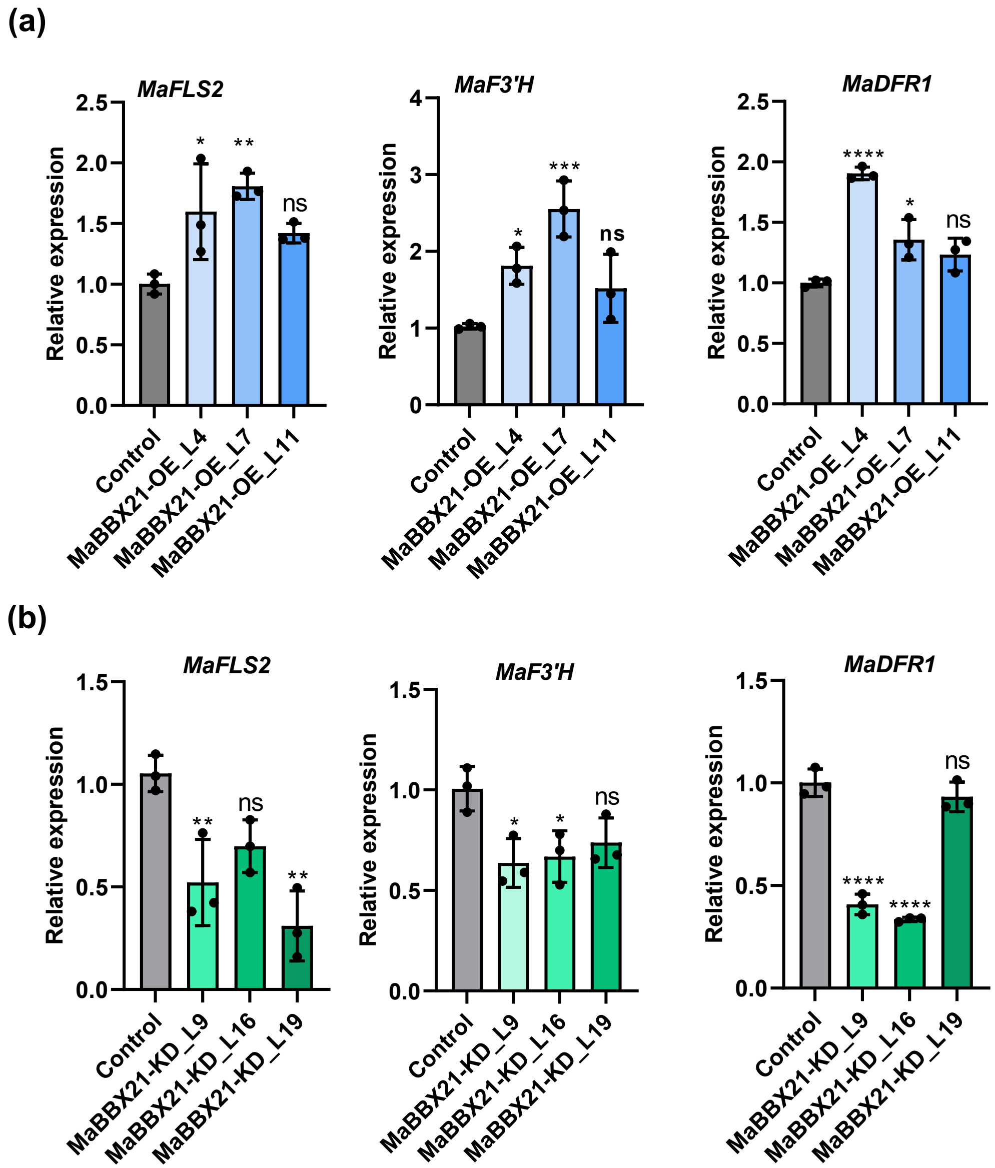
(a)
MaFLS2
Relative expression
Control
MaBBX21-OE_L4
MaBBX21-OE_L7
MaBBX21-OE_L11
*
**
ns
MaF3'H
***
MaDFR1
****
(b)
MaFLS2
MaBBX21-KD_L9
MaBBX21-KD_L16
MaBBX21-KD_L19
MaF3'H
MaDFR1

Figure S11

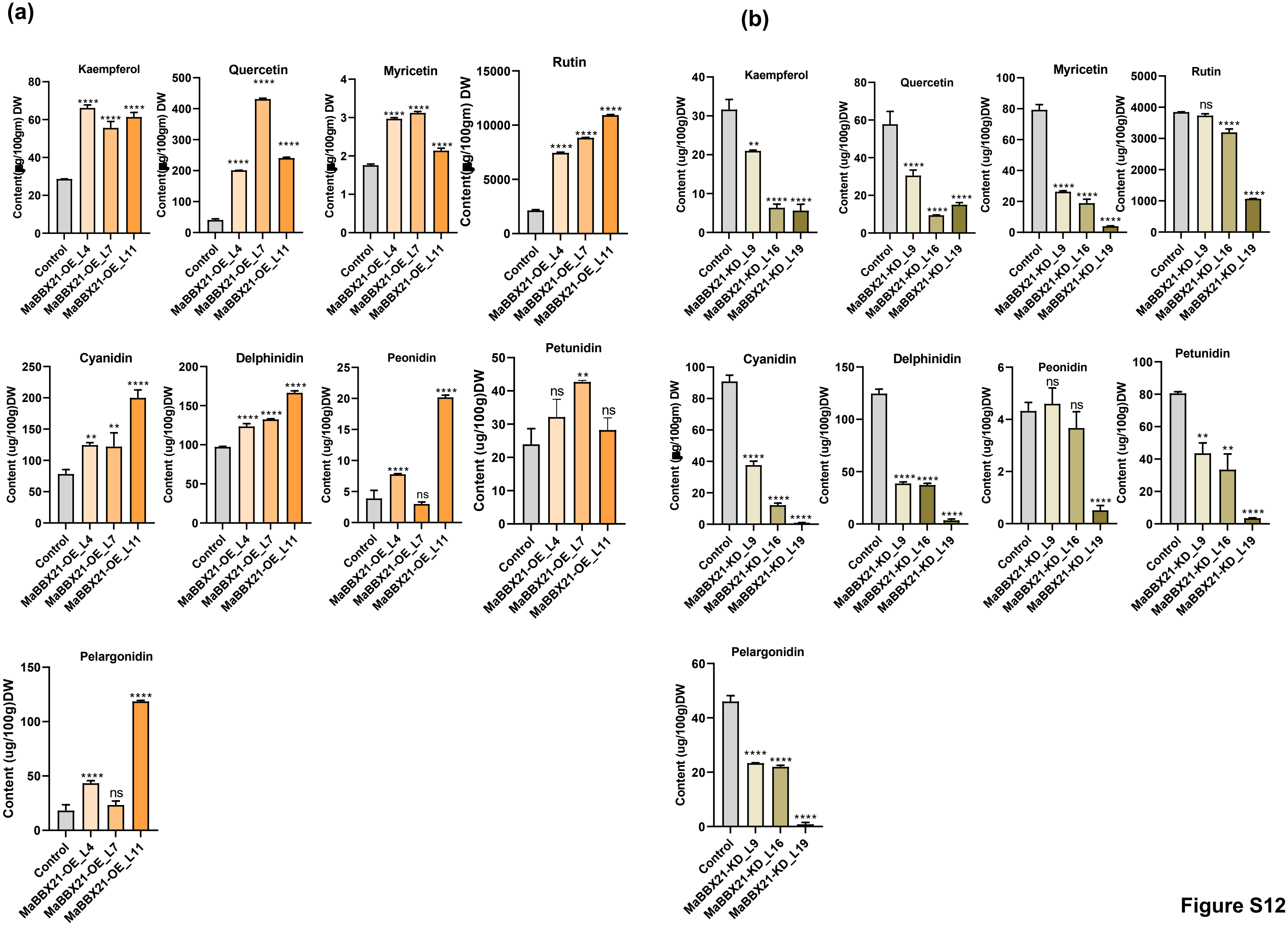
(a)
(b)
Kaempferol
Quercetin
Myricetin
Rutin
Cyanidin
Delphinidin
Peonidin
Petunidin
Pelargonidin
Content (ug/100gm) DW
Content (ug/100g)DW
Content (ug/100g) DW
Control
MaBBX21-OE_L4
MaBBX21-OE_L7
MaBBX21-OE_L11
MaBBX21-KD_L9
MaBBX21-KD_L16
MaBBX21-KD_L19
****
**
ns

Figure S12

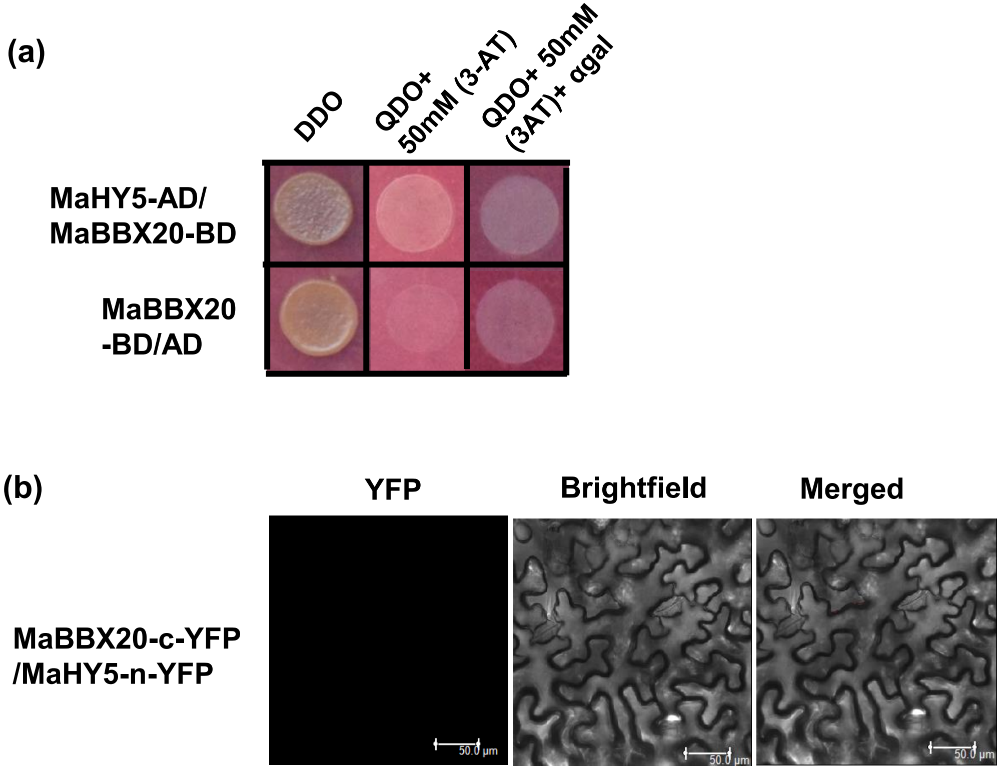

(a)
DDO
QDO+ 50mM (3-AT)
QDO+ 50mM (3AT)+ αgal
MaHY5-AD/ MaBBX20-BD
MaBBX20 -BD/AD
(b)
YFP
Brightfield
Merged
MaBBX20-c-YFP /MaHY5-n-YFP
50.0 µm


Figure S13

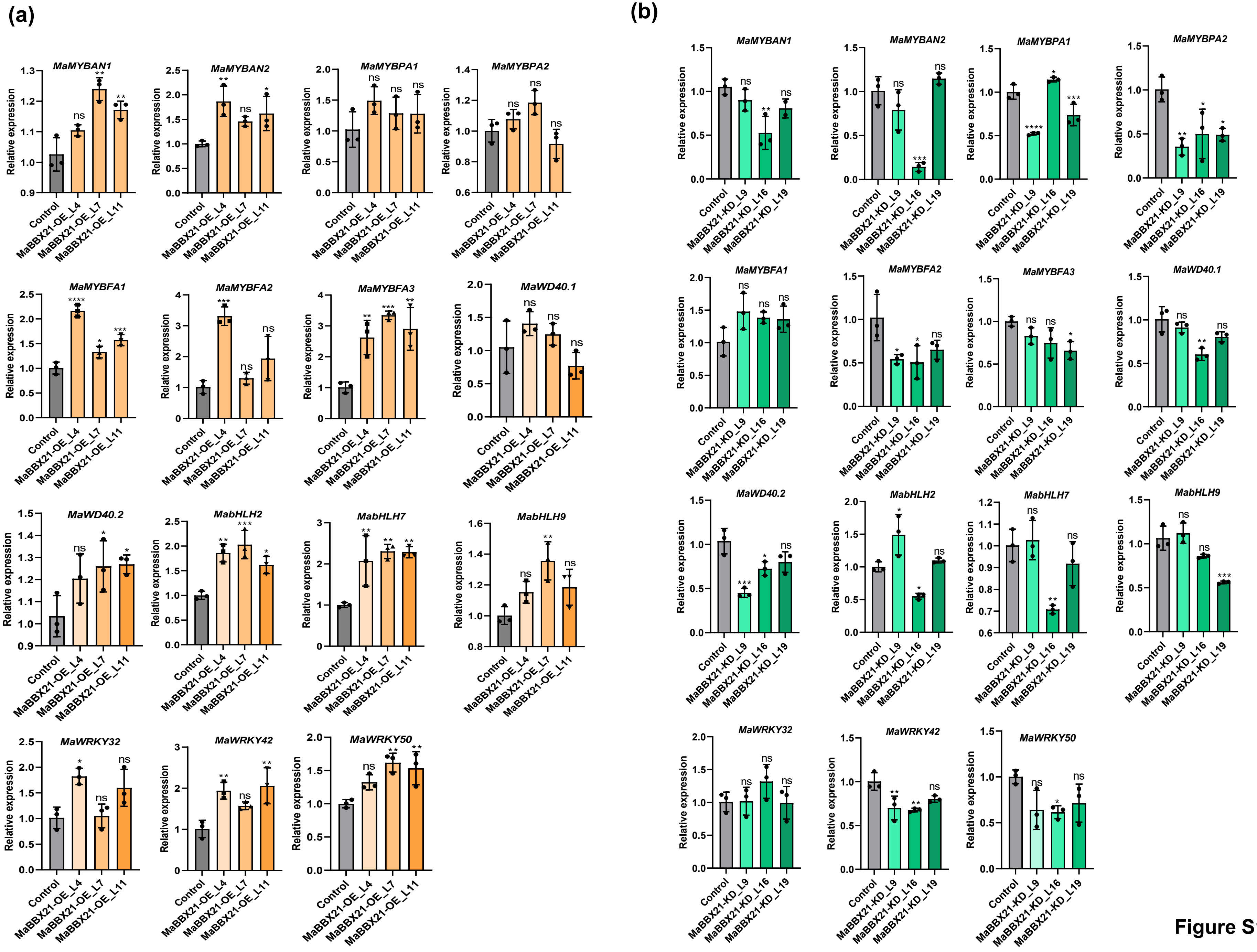
(a)
MaMYBAN1
MaMYBAN2
MaMYBPA1
MaMYBPA2
MaMYBFA1
MaMYBFA2
MaMYBFA3
MaWD40.1
MaWD40.2
MabHLH2
MabHLH7
MabHLH9
MaWRKY32
MaWRKY42
MaWRKY50
Relative expression
Control
MaBBX21-OE_L4
MaBBX21-OE_L7
MaBBX21-OE_L11
(b)
MaMYBAN1
MaMYBAN2
MaMYBPA1
MaMYBPA2
MaMYBFA1
MaMYBFA2
MaMYBFA3
MaWD40.1
MaWD40.2
MabHLH2
MabHLH7
MabHLH9
MaWRKY32
MaWRKY42
MaWRKY50
Relative expression
Control
MaBBX21-KD_L9
MaBBX21-KD_L16
MaBBX21-KD_L19

Figure S14

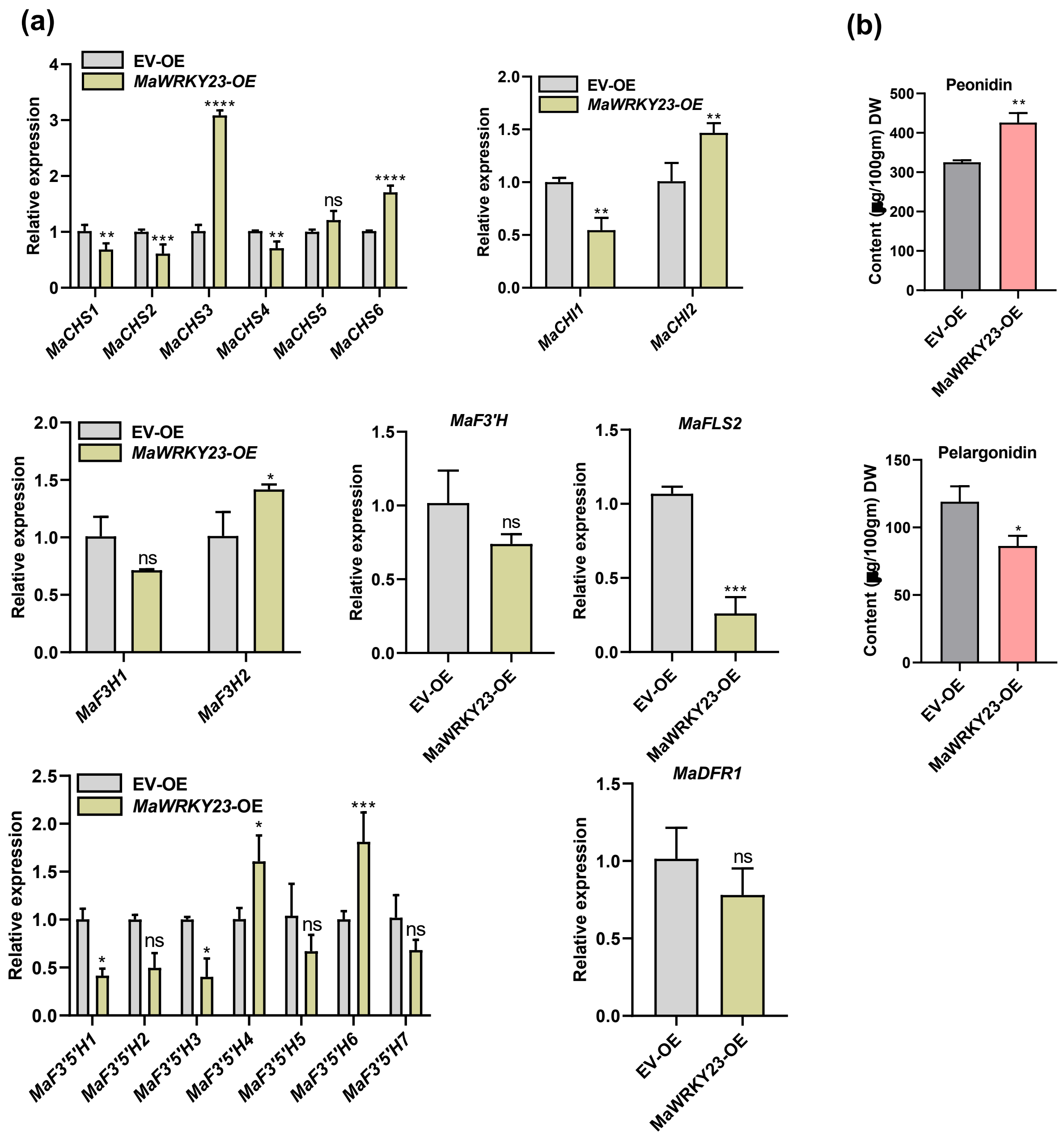
(a)
EV-OE
MaWRKY23-OE
Relative expression
MaCHS1
MaCHS2
MaCHS3
MaCHS4
MaCHS5
MaCHS6
MaCHI1
MaCHI2
MaF3H1
MaF3H2
MaF3'H
MaFLS2
MaDFR1
MaF3'5'H1
MaF3'5'H2
MaF3'5'H3
MaF3'5'H4
MaF3'5'H5
MaF3'5'H6
MaF3'5'H7
(b)
Peonidin
Pelargonidin
Content (μg/100gm) DW

Figure S15

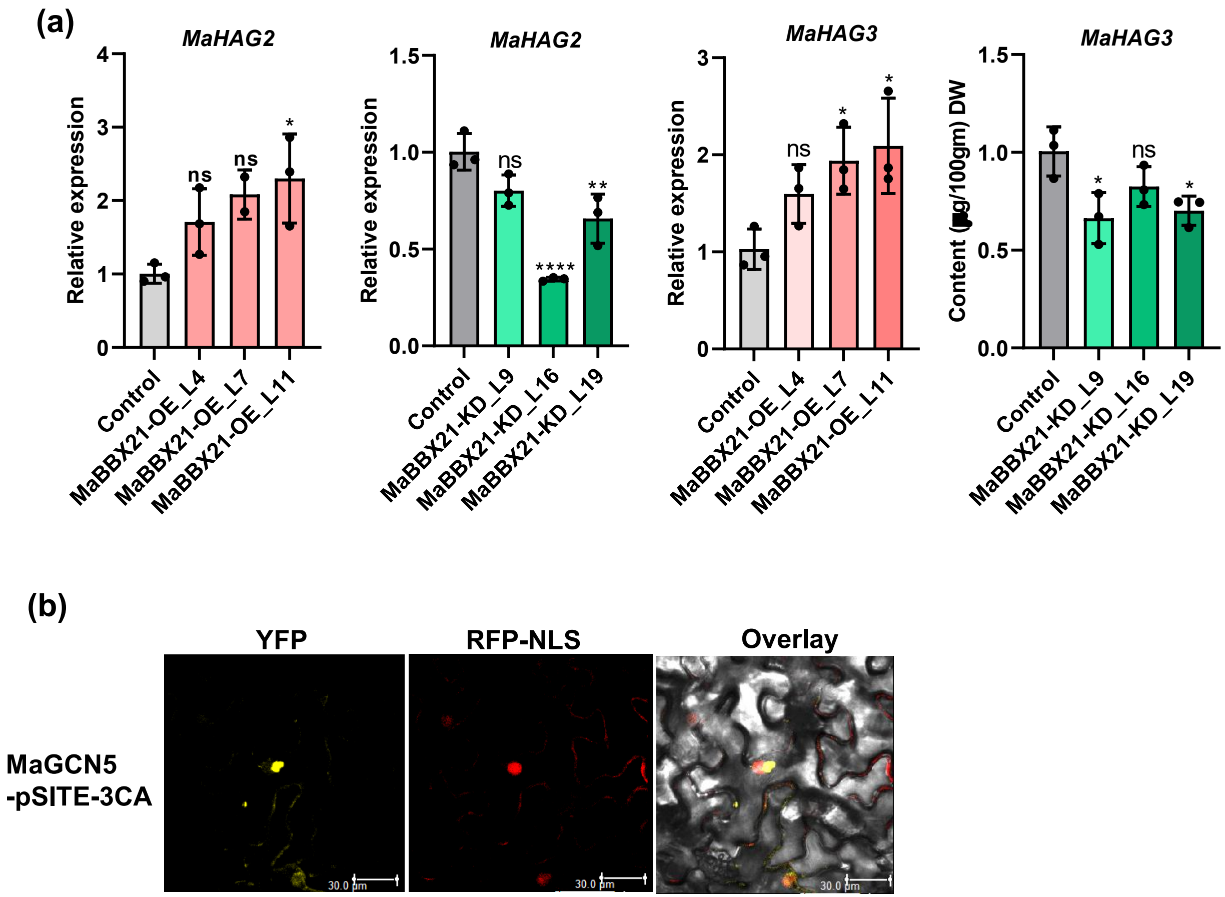
(a)
MaHAG2
Relative expression
ns
ns
*
Control
MaBBX21-OE_L4
MaBBX21-OE_L7
MaBBX21-OE_L11
MaHAG2
Relative expression
ns
****
**
Control
MaBBX21-KD_L9
MaBBX21-KD_L16
MaBBX21-KD_L19
MaHAG3
Relative expression
ns
*
*
MaHAG3
Content (µg/100gm) DW
*
ns
*
(b)
YFP
RFP-NLS
Overlay
MaGCN5
-pSITE-3CA
30.0 µm

Figure S16

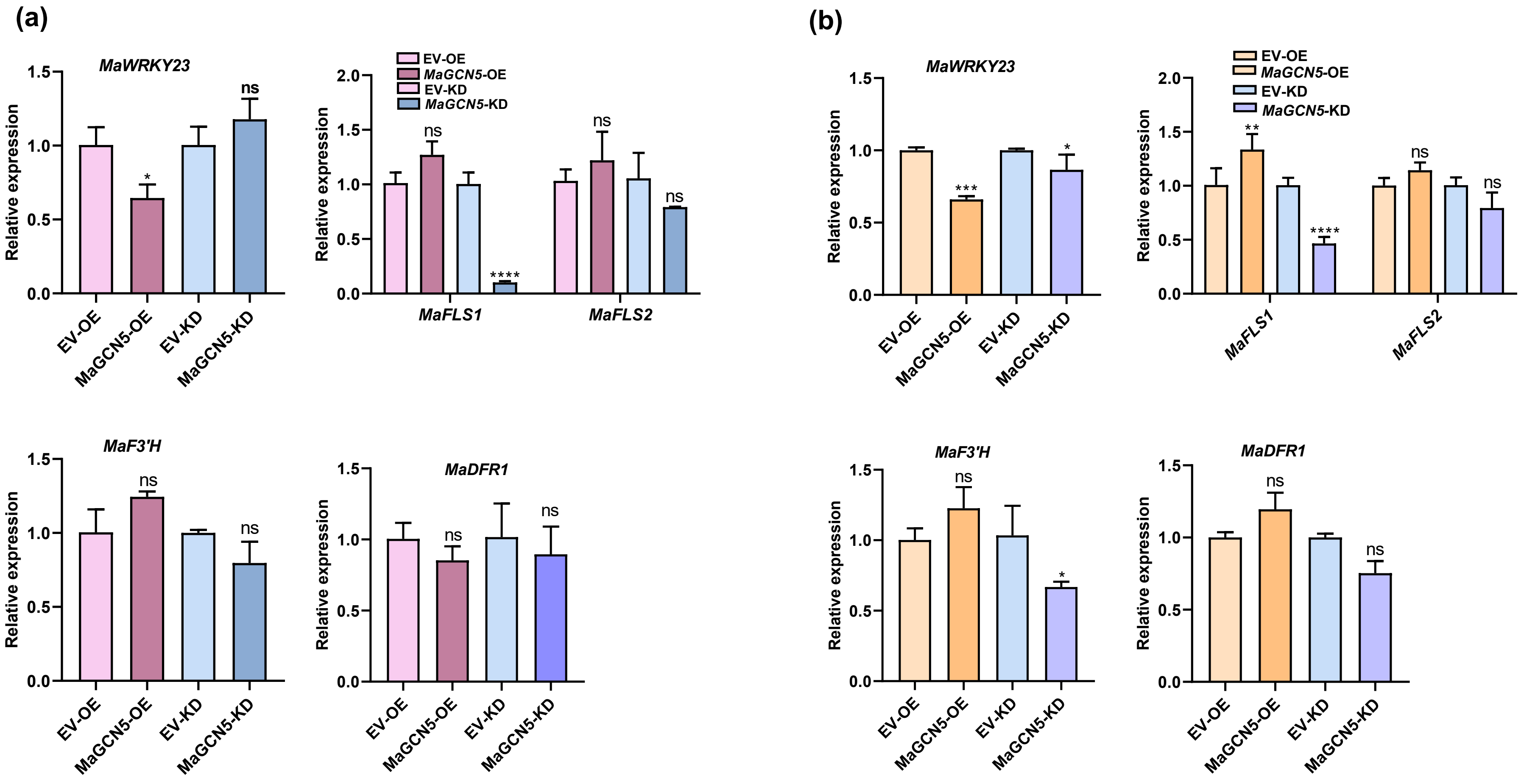
(a)
MaWRKY23
Relative expression
EV-OE
MaGCN5-OE
EV-KD
MaGCN5-KD
ns
MaFLS1
MaFLS2
MaF3'H
MaDFR1
(b)

Figure S17

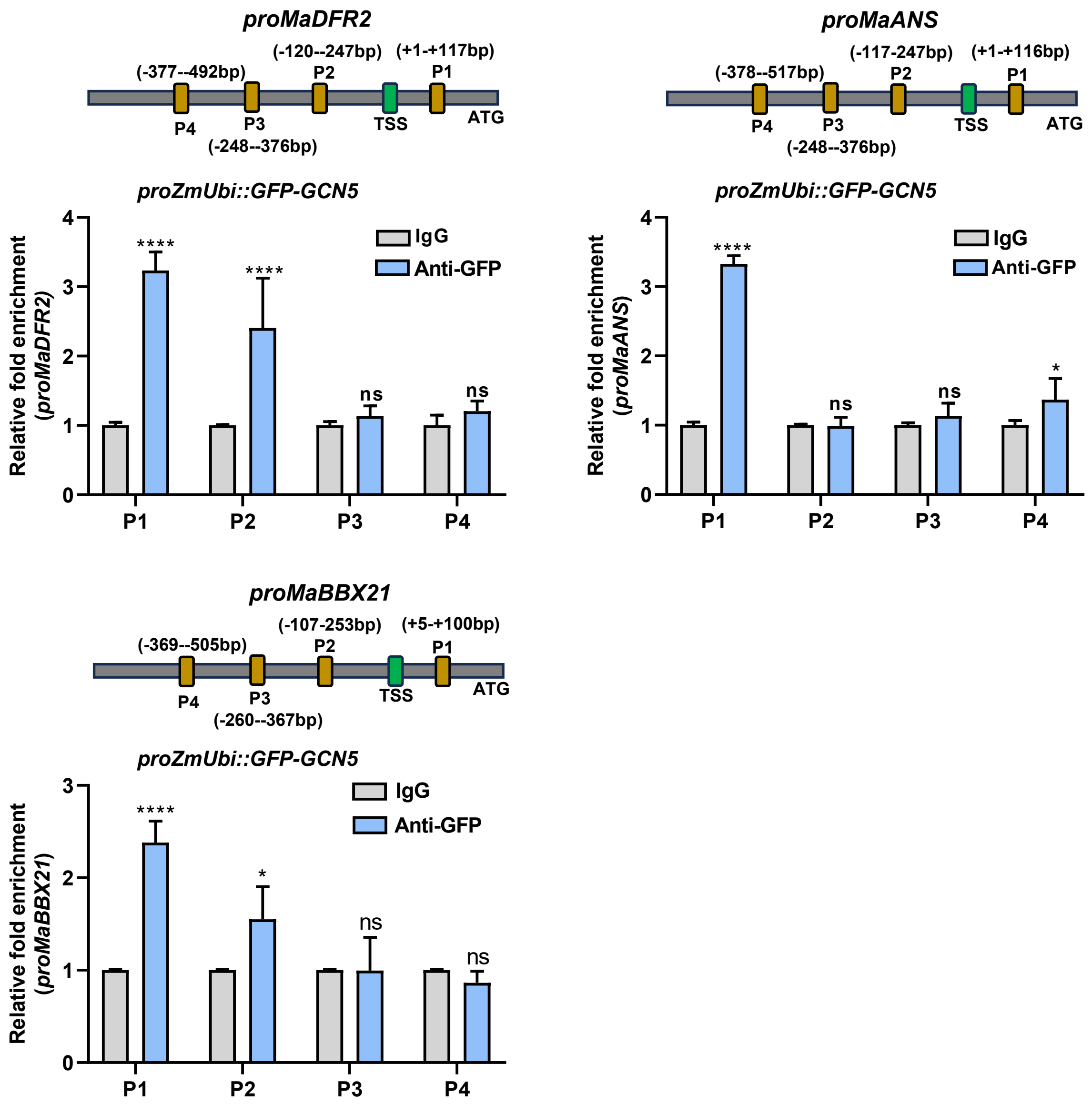
proMaDFR2
(-377--492bp)
(-120--247bp)
(+1-+117bp)
P2
P1
P4
P3
TSS
ATG
(-248--376bp)
proZmUbi::GFP-GCN5
Relative fold enrichment (proMaDFR2)
IgG
Anti-GFP
****
****
ns
ns
P1
P2
P3
P4
proMaANS
(-378--517bp)
(-117-247bp)
(+1-+116bp)
P2
P1
P4
P3
TSS
ATG
(-248--376bp)
proZmUbi::GFP-GCN5
Relative fold enrichment (proMaANS)
IgG
Anti-GFP
****
ns
ns
*
P1
P2
P3
P4
proMaBBX21
(-369--505bp)
(-107-253bp)
(+5-+100bp)
P2
P1
P4
P3
TSS
ATG
(-260--367bp)
proZmUbi::GFP-GCN5
Relative fold enrichment (proMaBBX21)
IgG
Anti-GFP
****
*
ns
ns
P1
P2
P3
P4

Figure S18

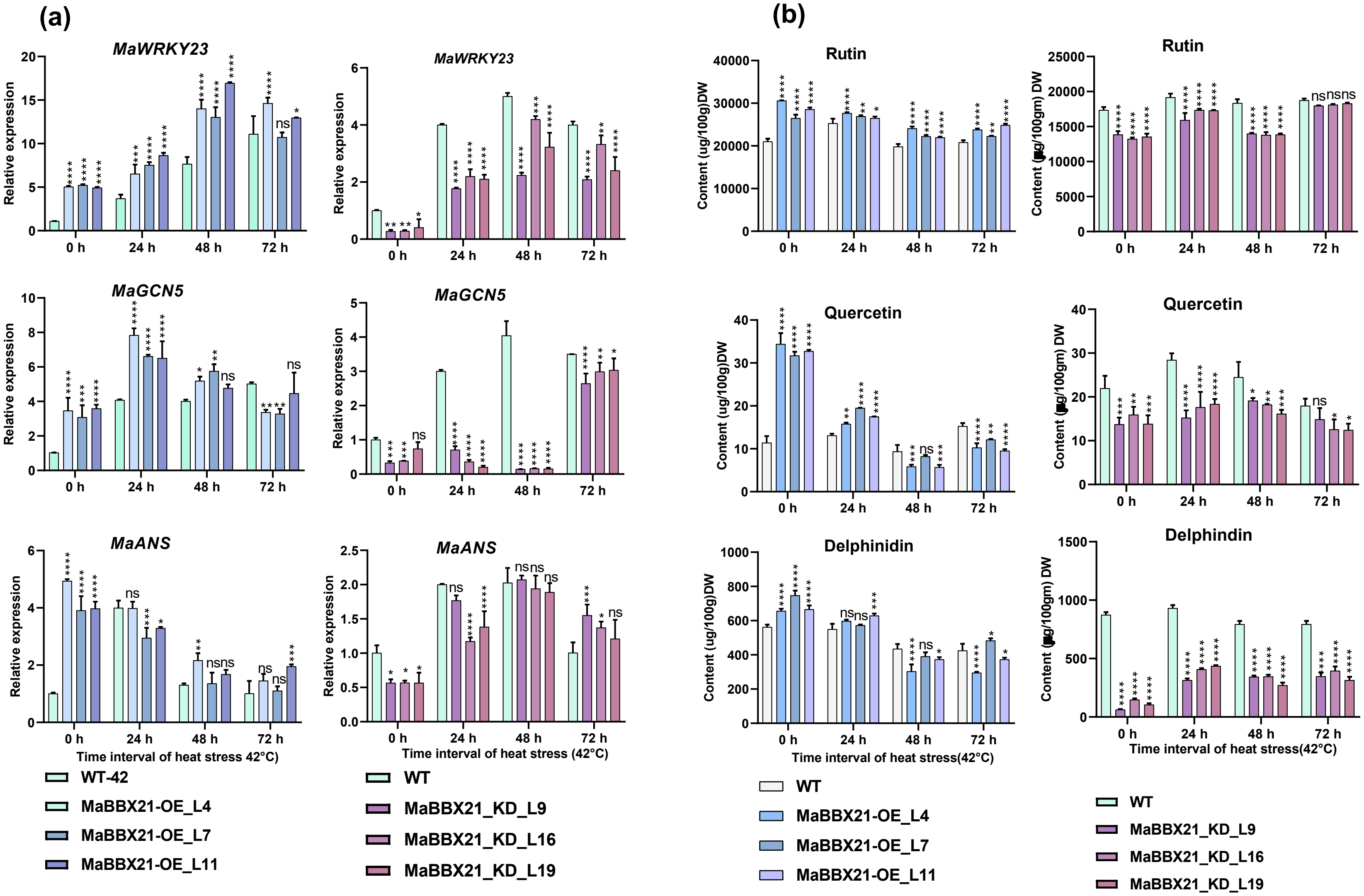

(a)
(b)
MaWRKY23
MaGCN5
MaANS
Relative expression
Rutin
Quercetin
Delphinidin
Delphindin
Content (ug/100g)DW
Time interval of heat stress 42°C)
Time interval of heat stress (42°C)
Time interval of heat stress(42°C)
WT-42
MaBBX21-OE_L4
MaBBX21-OE_L7
MaBBX21-OE_L11
WT
MaBBX21_KD_L9
MaBBX21_KD_L16
MaBBX21_KD_L19


Figure S19

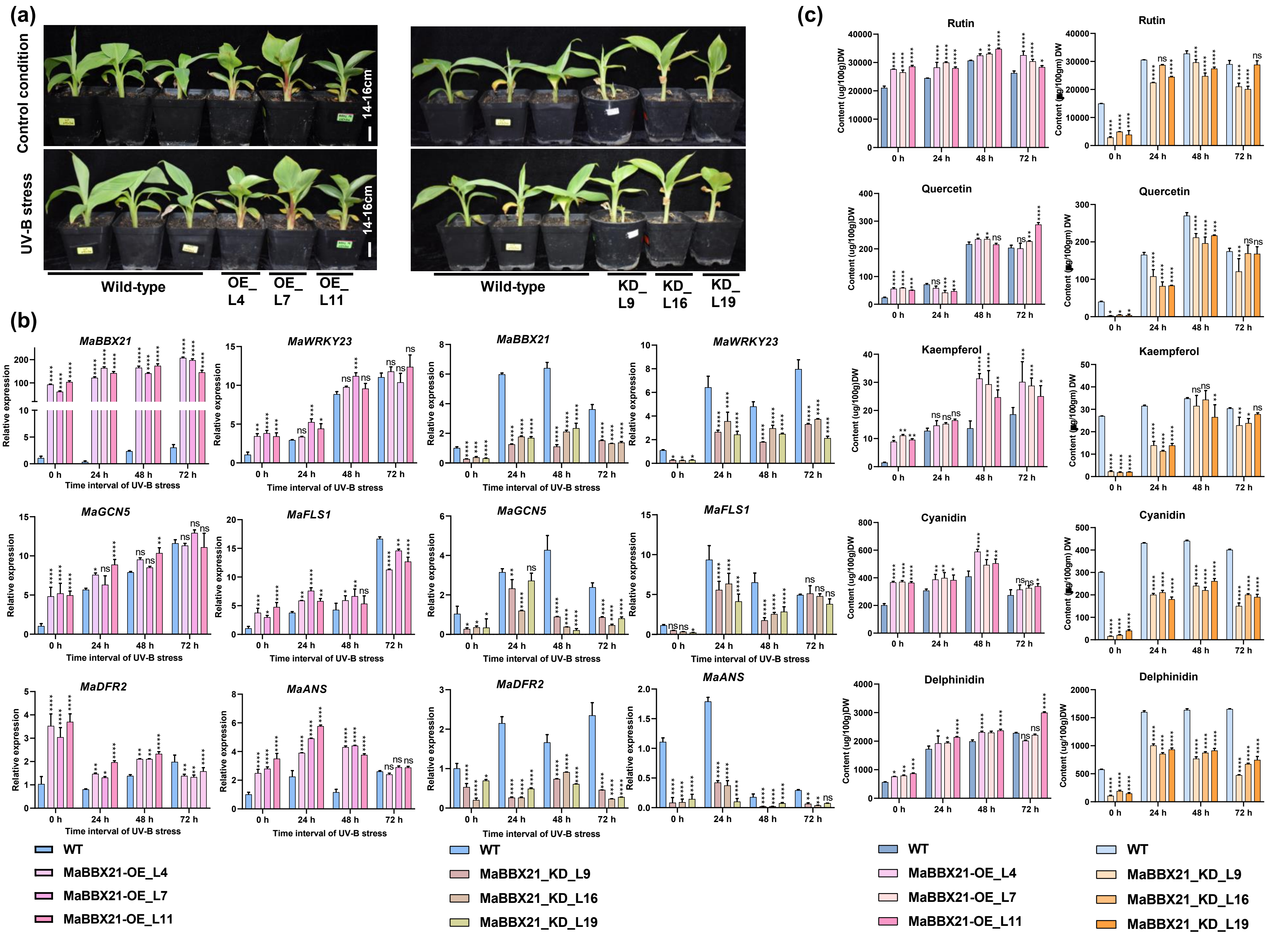
(a)
Control condition
UV-B stress
14-16cm
Wild-type
OE_L4
OE_L7
OE_L11
KD_L9
KD_L16
KD_L19
(b)
MaBBX21
MaWRKY23
MaGCN5
MaFLS1
MaDFR2
MaANS
Relative expression
Time interval of UV-B stress
0 h
24 h
48 h
72 h
WT
MaBBX21-OE_L4
MaBBX21-OE_L7
MaBBX21-OE_L11
MaBBX21_KD_L9
MaBBX21_KD_L16
MaBBX21_KD_L19
(c)
Rutin
Quercetin
Kaempferol
Cyanidin
Delphinidin
Content (ug/100g)DW
Content (ug/100gm) DW

Figure S20

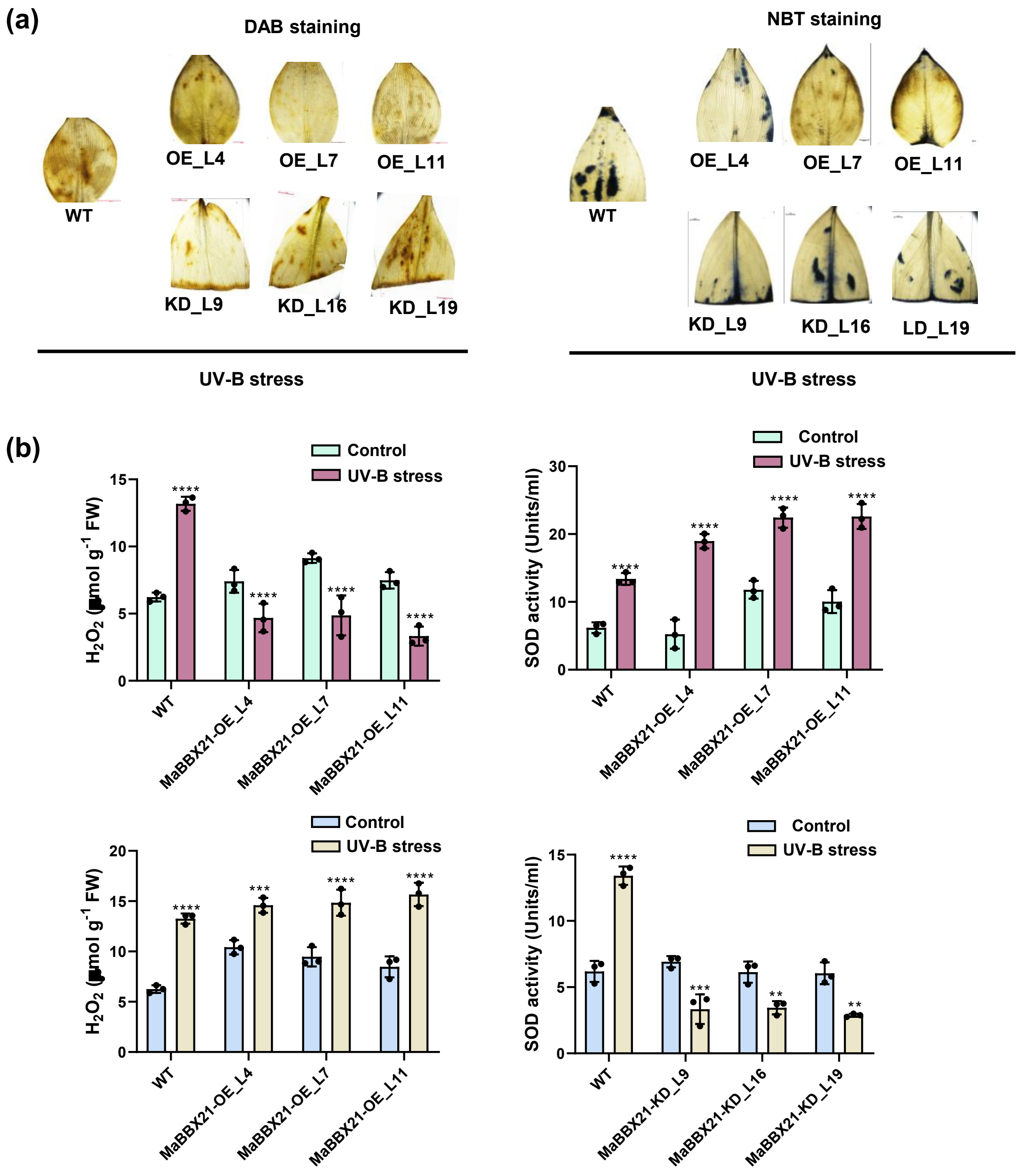

(a)
DAB staining
WT
OE_L4
OE_L7
OE_L11
KD_L9
KD_L16
KD_L19
UV-B stress
NBT staining
WT
OE_L4
OE_L7
OE_L11
KD_L9
KD_L16
LD_L19
UV-B stress
(b)
Control
UV-B stress
H2O2 (µmol g-1 FW)
SOD activity (Units/ml)
WT
MaBBX21-OE_L4
MaBBX21-OE_L7
MaBBX21-OE_L11
MaBBX21-KD_L9
MaBBX21-KD_L16
MaBBX21-KD_L19


Figure S21
